# Evolution of irreversible differentiation under stage-dependent cell differentiation

**DOI:** 10.1101/2023.05.04.539351

**Authors:** Yuanxiao Gao, Román Zapién-Campos, Yuriy Pichugin, Arne Traulsen

**Affiliations:** School of Mathematics and Data Science, Shaanxi University of Science and Technology, 710021 Xi’an, China; Max Planck Institute for Evolutionary Biology, August-Thienemann-Str. 2, 24306 Plön, Germany; Centre for Life’s Origins and Evolution, University College London, WC1 6BT, London, UK; Department of Ecology and Evolutionary Biology, Princeton University, Princeton, NJ, USA

**Author notes:** Corresponding author: Yuanxiao Gao.

## Abstract

The specialization of cells is a hallmark of complex multicellularity. Cell differentiation enables the emergence of specialized cell types that carry out separate functions previously executed by a multifunctional ancestor cell. One view is that initial cell differentiation occurred randomly, especially for genetically identical cells, exposed to the same life history environment. How such a change in differentiation probabilities can affect the evolution of differentiation patterns is still unclear. We develop a theoretical model to investigate the effect of stage-dependent cell differentiation – cells change their developmental trajectories during a single round of development via cell divisions – on the evolution of optimal differentiation patterns. We found that irreversible differentiation – a cell type gradually losing its differentiation capability to produce other cell types – is more favored under stage-dependent than stage-independent cell differentiation in relatively small organisms with limited differentiation probability variations. Furthermore, we discovered that irreversible differentiation of germ cells, which is the gradual loss of germ cells’ ability to differentiate, is a prominent pattern among irreversible differentiation patterns under stage-dependent cell differentiation. In addition, large variations in differentiation probabilities prohibit irreversible differentiation from being the optimal differentiation pattern.

**Author summary:** The differentiation of cells into different branches is a characteristic feature of multicellular organisms. To understand its origin, the mechanism of division of labour was proposed, where cells are specialized at distinct tasks. In previous models, a cell type is usually assumed to produce another cell type with a fixed probability which is referred to as stage-independent differentiation. However, it has been argued that cell differentiation is a dynamic process in which cells possess changing differentiation capabilities during the different stages of an organism’s development. Stage-dependent differentiation exhibits more diverse patterns of development than differentiation with fixed probabilities, thus it can lead to novel targets of selection. How does stage-dependent differentiation impact the evolution of optimal differentiation patterns compared with stage-independent one? To address this question, we built a stage-dependent cell differentiation model and classified differentiation patterns based on the cells’ differentiation capability in their last cell division. We investigate how stage-dependent differentiation probabilities impact the evolution of the optimal differentiation pattern, which acts on the fitness of an organism. As we take the growth rate as a proxy of an organism’s fitness, we seek the “optimal strategy” that leads to the fastest growth. Our numerical results show that irreversible differentiation which gradually loses its differentiation capability, is favored over stage-independent differentiation in small organisms. Meanwhile, irreversible differentiation won’t be optimal when there are no constraints on the changes of stage-dependent differentiation probabilities between successive cell divisions.

## Introduction

The evolution of multicellularity has been viewed as the major evolutionary transition for the evolution of life on earth Maynard Smith and Szathmáry [1995], Szathmáry and Smith [1995], Ratcliff et al. [2015], Sebe-Pedros et al. [2017], Márquez-Zacarías et al. [2021b]. One important aspect of this is cell differentiation into different cell types. Cooperation and division of labor between these cells have been widely investigated in the evolution of multi-cellularity Ratcliff et al. [2012], Hammerschmidt et al. [2014], West et al. [2015], Gao et al. [2019], Rose et al. [2020]. Multicellular organisms, especially large ones, possess different cell types to perform diverse functions Carroll [2001], McCarthy and Enquist [2005], Arendt [2008]. It is widely accepted that multicellular life has evolved from unicellular ancestors Mikhailov et al. [2009], Claessen et al. [2014]. Division of labour in organisms enables a diversity of cell types, leading unicellular organisms to form increasingly larger and more complex organizations. Differentiated cells perform distinct functions in varying conditions and can in this way increase an organism’s reproductive fitness. For example, cell differentiation occurs under adverse environmental conditions to increase an organism’s survival chance, such as cyanobacteria differentiating nitrogen-fixing heterocysts to use *N*_2_ when combined-nitrogen is insufficient Gallon [1992], *Saccharomyces cerevisiae* producing cells with different apoptosis likelihood under gravitational selection Ratcliff et al. [2012] or *Myxococcus xanthus* producing a new cell type under starvation Claessen et al. [2014].

Several mechanisms have been proposed to understand cell differentiation and phenotypic variation, from the perspective of gene expression, mutations, epigenetics, and the environment Extavour and Akam [2003], Arendt [2008], Mikhailov et al. [2009], West and Cooper [2016], Arendt et al. [2016], Brunet and King [2017], Márquez-Zacarías et al. [2021b], Huang et al. [2024]. These mechanisms are complementary, and thus, more than one mechanism could act during the evolution of cell differentiation West and Cooper [2016], Brunet and King [2017]. These mechanisms usually assume that multifunctional and unicellular ancestors differentiate into specialized cells to carry out segregated functions, even when cells are genetically identical and have been exposed to an identical environment. It has been shown that cells differentiate depending on the development states of an organism. For example, one out of successive 10 to 15 vegetative cells differentiate into a new cell type, heterocyst, in filamentous cyanobacteria *Anabaena sp*.PCC 7120 Flores and Herrero [2010]; *Volvox* differentiates into two cell types at its 6th round of division in its whole 11 ∼ 12 rounds of cell divisions Matt and Umen [2016]. Moreover, in closed related species of *Volvox* family, it has been found that the observed stable differentiation patterns are highly likely the evolution consequences of originally randomly happened cell differentiation. For instance, smaller species *Gonium* have identical cells, whereas intermediate-sized species *Volvox aureus* and *Volvox gigas* have partial germ-soma differentiation, whereas *Volvox carteri* and *Volvox obversus* have complete germ-soma differentiation Matt and Umen [2016]. How originally occurred state-dependent cell differentiation in an organism shapes the evolution of cell differentiation patterns is still unclear.

Studies of cell differentiation have mainly focused on the optimal condition, where mature cells of an organism allocate their resources to different tasks Michod [2007], Willensdorfer [2009], Gavrilets [2010], Rossetti et al. [2010], Rueffler et al. [2012], Ispolatov et al. [2012], Solari et al. [2013], Goldsby et al. [2014], Cooper and West [2018], Liu et al. [2021], Cooper et al. [2021, 2022]. Essentially, they are focused on the proportion of each cell type in an organism, instead of the stochastic developmental process of each cell type during an organism’s growth. Cells capable of switching to another cell type have not been in the focus yet. Some authors considered cell differentiation abilities, but only in one cell type while other cell types were terminally differentiated types (without division ability) Willensdorfer [2009], Rossetti et al. [2010], Solari et al. [2013]. Rodrigues et al. considered cell differentiation ability as an evolving trait, but the trait was coupled with varying cell division rates and organisms were constrained to filament form Rodrigues et al. [2012]. More recently, Cooper et al. introduced a random specialization model, but the random process only impacts the final fractions of different cell types rather than the internal organization of task allocation of cells during an organism’s growth Cooper et al. [2022]. Gavrilets and Gao et al. considered cell differentiation between cell types, but the differentiation probabilities are assumed to be fixed rather than stochastic Gavrilets [2010], Gao et al. [2021]. So far, little is known about the effects of stage-dependent differentiation probabilities on the evolution of cell differentiation patterns, such as irreversible or reversible.

In this study, based on our previous work Gao et al. [2021], we develop a theoretical model to investigate the effect of cell differentiation with stage-dependent differentiation probabilities on the evolution of optimal differentiation patterns. Stage-dependent cell differentiation refers to the capability of cells having different cell differentiation probabilities between any two successive cell divisions. Comparatively, stage-independent differentiation only allows a cell type to have a fixed cell differentiation probability across cell divisions Gao et al. [2021]. Inspired by the cells’ division of labour of *Volvox*, where germ cells are responsible for reproduction and somatic cells are responsible for viability Matt and Umen [2016], we consider two cell types in an organism: germ-like cells and soma-like cells. We use the expected offspring number of an organism i.e. growth rate as a proxy of an organism’s fitness because it is the simplest direct criterion Parker and Smith [1990]. We assume that an organism grows by cell divisions which further depends on the fraction of soma-like cells and transition probabilities between cell types. Different stage-dependent strategies compete to maximize the organism’s fitness. We numerically calculate organisms’ growth rates under different parameters and compare the evolutionary differences of optimal strategies under stage-dependent and stage-independent cell differentiation. Intuitively, reversible differentiation instead of irreversible differentiation under stage-dependent differentiation will be selected especially when cost is low, because reversible differentiation can “recycle” soma-type cells for reproduction. However, we found that stage-dependent differentiation favors irreversible differentiation more than stage-independent differentiation even without costs in small organisms.

## Model and methods

We designed a life cycle model for organisms with stage-dependent cell differentiation compared with previous work investigated under stage-independent cell differentiation Gao et al. [2021]. As we are focused on the formation process of differentiation patterns, we consider two intermediate cell types rather than specific cell types in an organism: germ-like and soma-like, which is inspired by the partial differentiation cell types in genus *Pandorina*, a closed genus of *Volvox* Matt and Umen [2016]. Here, the two cell types are allowed to differentiate into each other, and we investigate the possible differentiation process along with cell divisions, which we refer to as differentiation strategies. Different strategies lead organisms to different developmental trajectories and fitness. In the model, an organism’s growth rate is a fitness proxy as it is the simplest direct way to measure organisms’ fitness Parker and Smith [1990]. Next, we introduce the definition of differentiation strategies. We assume that each organism starts with a single germ-like cell, see Fig 1A. Cells divide synchronously, each cell producing two daughter cells at a time. After the *i*th cell division, organisms have 2^*i*^ cells in total. Organisms grow and mature until they reach a maturity size 2^*n*^, where *n* is the maximal cell division of organisms. Each germ-like cell is released from a mature organism as offspring to start a new life cycle. All soma-like cells in a mature organism die. For each division, cells have a set of probabilities to produce daughter cells of a certain type. Here, 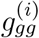 is the probability of a germ-like cell producing two germ-like cells at the *i*th cell division. The probabilities 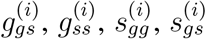 and 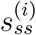 are defined in a similar manner, where we have 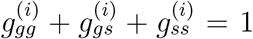 and 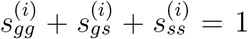 for each growth stage *i, i* = 1, 2, …, *n*. We denote 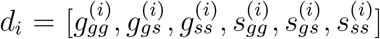 as the cell differentiation probabilities in the *i*th cell division. In addition, 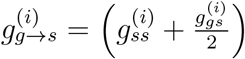 and 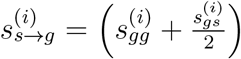 are referred to as transition probabilities, *i* = 1, 2, …, *n*. The cell differentiation probabilities across the successive *n* rounds of cell divisions of an organism can be expressed in matrix form as

**Figure 1:**
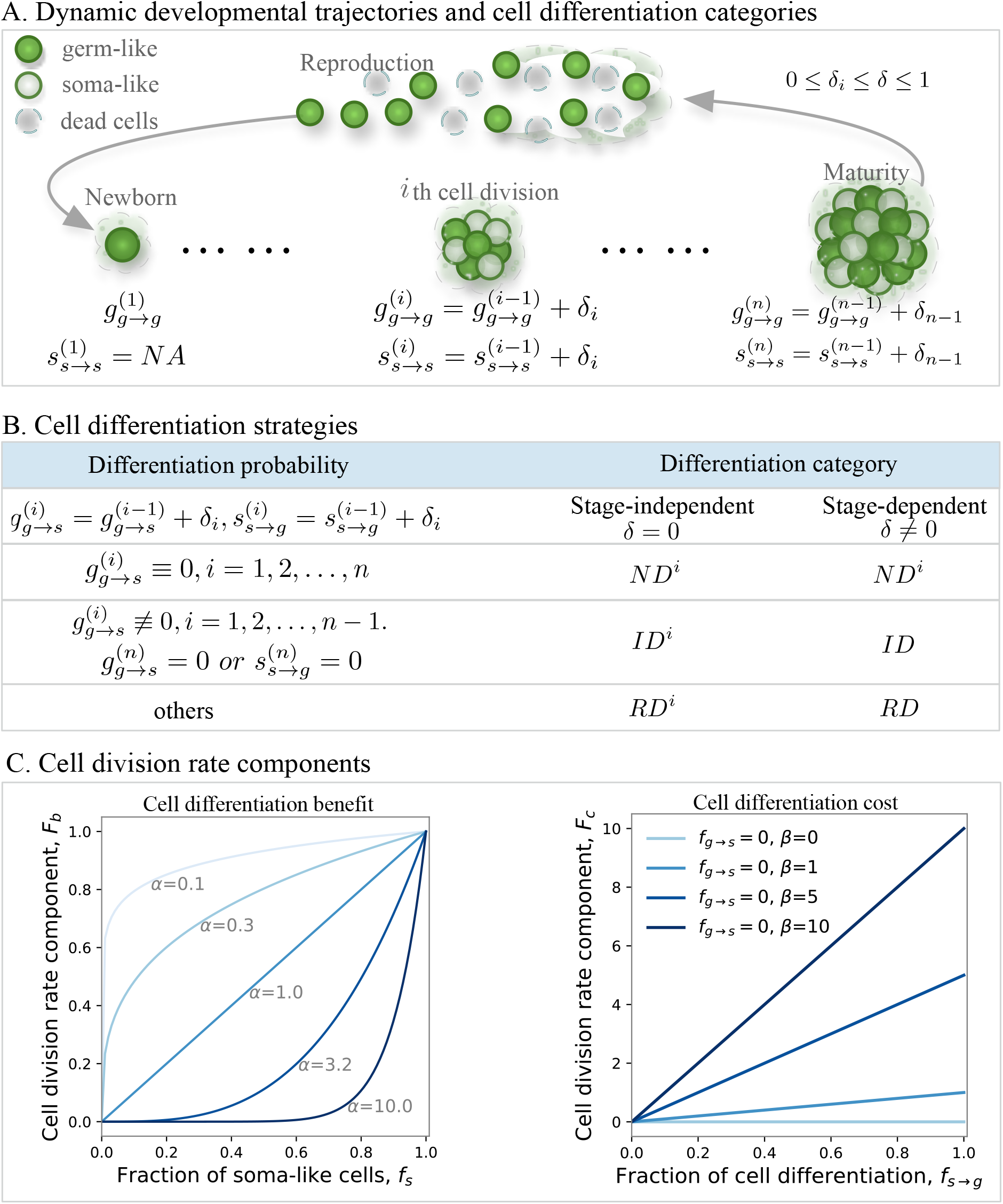
Illustration of the stage-dependent cell differentiation, differentiation strategies, and cell division rate components. **A**. Schematic of an organism’s life cycles. Organisms start from single germ-like cells and undergo *n* synchronous cell divisions before reproduction. For newborn organisms, the cell differentiation probability for soma-like cells is irrelevant as there are no soma-like cells. Cell differentiation probabilities can change from the (*i* − 1)th cell division to the *i*th cell division by a small quantity *δ*_*i*_ (0 ≤ *δ*_*i*_ ≤ 1 and *i* = 1, 2, …, *n*). *δ* is the maximum change between successive cell differentiation probabilities i.e. 0 ≤ *δ*_*i*_ ≤ *δ* ≤ 1, *i* = 1, …, *n*. **B**. Cell differentiation strategy classification. Based on the cell differentiation probabilities at the last cell division, we classify cell differentiation into three categories: non-differentiation *ND*, reversible differentiation *RD*, and irreversible differentiation *ID*. The upper script *i* means the strategy is stage-independent i.e. *δ* = 0. For *ND*, since 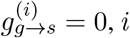, *i* = 1, …, *n*, thus *ND* equals *ND*^*i*^. **C**. Cell division rate components. The left panel shows the benefits of cell differentiation. We assume that the cell division rate increases with the fraction of soma-like cells *f*_*s*_. For the associated benefit, we assume *F*_*b*_ = *b*(*f*_*s*_)^*α*^, where the shape of the function is controlled by *α*. The right panel shows the costs of cell differentiation. We assume that the cell division rate decreases with the fraction of cell divisions that turn a soma-like cell into a germ-like cell and vice versa. For the associated cost, we assume *F*_*c*_ = *c*(*f*_*g*→*s*_ + *βf*_*s*→*g*_). Here, we show the values of *F*_*c*_ with varying *f*_*s*→*g*_ and *β* by setting *f*_*g*→*s*_ = 0 (Parameters: *b* = 1 in the left panel and *c* = 1 in the right panel).

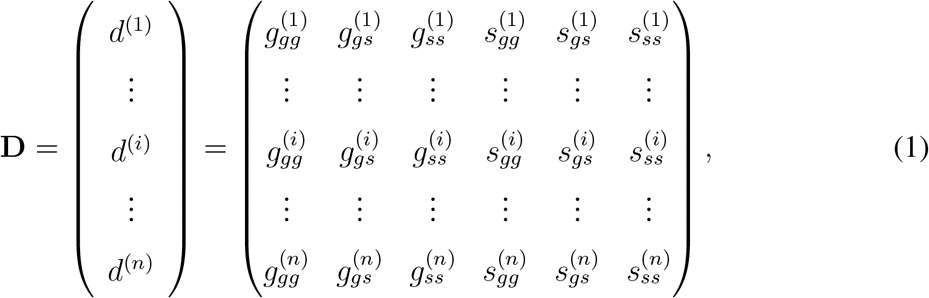

where the *i*th row of the matrix contains the cell differentiation probabilities in the *i*th cell division. We call **D** stage-dependent cell differentiation, as the probabilities can change between different division stages. We assume that this change is not larger than *δ*, e.g 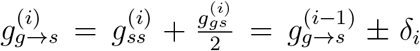 with *δ*_*i*_ ≤ *δ* sufficiently small such that all probabilities remain well defined. If *δ* = 0 for *i* = 1, 2, …, *n*, then **D** is a stage-independent cell differentiation strategy, where the same type of cells follow a fixed set of probabilities to produce daughter cells at each division. We should note that the stage-independent cell differentiation is also defined by the values of 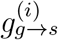 and 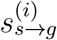 but which don’t change across *i*. Different cell differentiation strategies lead to different differentiation degrees. For example, if a strategy has 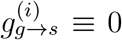 for *i* = 1, 2, …, *n*, then organisms have no cell differentiation. If a strategy has 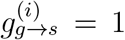 and 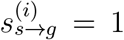, then organisms have maximal degree of cell differentiation. Stage-dependent differentiation allows many different trajectories. To distinguish them and focus on differentiation patterns, we consider the probabilities in the last division (Fig 1B). If 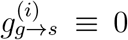 for *i* = 1, 2, …, *n*, then we call the differentiation non-differentiation *ND*. Otherwise, if germ-like cells differentiate soma-like cells at least one time before final cell division, i.e. 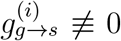, *i* = 1, 2, …, *n* − 1, but with NO differentiation for either cell type at last division i.e. 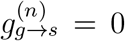 or 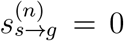, then we call it irreversible differentiation *ID*. Strategy *ID* captures the process by which cells gradually lose their differentiation capabilities. The rest differentiation is called reversible differentiation *RD*. We should stress the limitation of this classification, in which different strategies could lead to a similar development trajectory, especially in large organisms. Nevertheless, the classification is a simple way that distinguish different differentiation patterns. For convenience, we use the upper script *i* to show the stage-independent strategies where cells have fixed cell differentiation probabilities. From the definition of *ND*, we know it is a stage-independent cell differentiation, thus we use *ND*^*i*^ to denote it afterward. For strategy *ID*^*i*^, only soma-like cells can possess irreversibility i.e. *s*_*s*→*g*_ = 0 Gao et al. [2021]. In this work, the acronyms of differentiation strategies are stage-dependent unless otherwise stated.

We assume that cell differentiation impacts an organism’s growth Gao et al. [2021]. The effects of cell differentiation are further decomposed into cell differentiation benefits and costs on growth. We assume differentiated soma-like cells are beneficial and increase an organism’s growth. The assumption is based on the division of labour in *Volvox*, where somatic cells are responsible for viability and germ cells are responsible for reproduction Matt and Umen [2016]. Cell differentiation between germ-like cells and soma-like cells is costly and decreases growth. A direct impact of differentiation is the decreased number of offspring as the resource that could be used for reproduction to convert to newly typed somatic cells. Organisms grow faster with higher cell division rates and vice versa. Specifically, *r*^(*i*)^ represents the growth rate in the *i*th cell division and is determined by two components

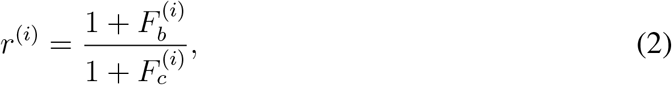

where 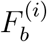 and 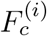 are the effects of cell differentiation benefit and cell differentiation cost in the *i*th cell division, *i* = 1, 2, …, *n. F*_*b*_ is a function of the fraction of soma-like cells *f*_*s*_,

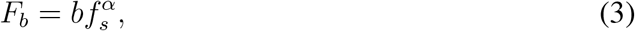

where the *b* is the benefit scale, *b* ≥ 0. *α* controls the shape of the function, see Fig 1C. *F*_*c*_ is a function of the fraction of cell differentiation between germ-like cell and soma-like cell *f*_*g*→*s*_ and *f*_*s*→*g*_,

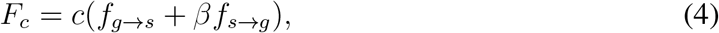

where *c* is the cost scale, *c* ≥ 0. *β* measures the relative cost of differentiation from soma-like cell to germ-like cell, see Fig 1C. The fractions of cell differentiation in the *i*th cell division are

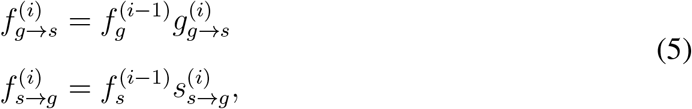

where 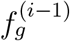 and 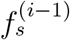 are the fraction of germ-like cell and soma-like cell after the (*i* − 1)th cell division, respectively. Note that 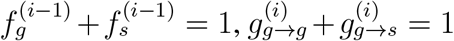, and 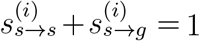, *i* = 1, 2, …, *n*. Specifically, 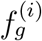and 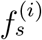are calculated by using transition probabilities, see Eq (11) in S1 Appendix. Taking Eq (2), Eq (3), Eq (4) and Eq (5) together, we have

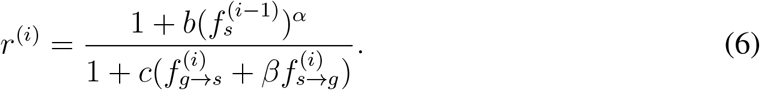

After the (*i* − 1)th cell division, the waiting time before the *i*th cell division occurring *t*^(*i*)^ follows the exponential distribution 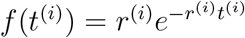, *i* = 1, 2, …, *n*. Thus the expected waiting time from the (*i*−1)th to the *i*th cell division is 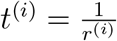 Allen [2010]. The expected growth time of organisms with *n* rounds of cell divisions is

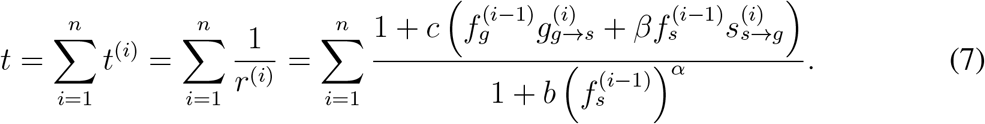

We consider a density-independent population, see Ress et al. [2022] for a discussion when density dependence is relevant in a related model. The growth rate of an organism only depends on the number of offspring and their growth time. As organisms divide synchronously, the number of offspring that an organism produces during its life is 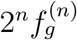 The expected number of offspring per unit time of an organism captures the effect of a given strategy on organisms. Therefore, organisms grow exponentially and an organism’s growth rate can be approximated by

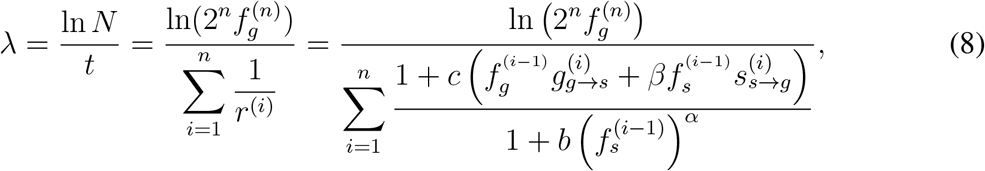

where *n* is the number of cell divisions an organism undergoes before maturity. 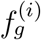and 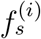 are fractions of germ-like cell and soma-like cell after the *i*th cell division. Here, 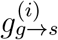 and 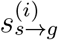 are the transition probabilities between germ-like cell and soma-like cell in the *i*th cell division (1 ≤ *i* ≤ *n*), see the S1 Appendix. We provide the calculation details of the growth rate in S1 Appendix and S2 Appendix.

It should be noted that the growth rate calculated here is not the exact growth rate for each realization. As each strategy in the model is stochastic, each strategy has different potential developmental trajectories. Therefore the growth rate of an organism under a strategy is a random variable depending on the probability of each trajectory that an organism can develop. Thus, the growth rate calculated via Eq (8) is an approximation of the mean growth rate. We test the robustness of the approximation in S3 Appendix. Our results show that the approximation is consistent with the mean growth rate of an organism. This model generalizes our previous study of stage-independent developmental trajectories using individual-based simulations Gao et al. [2021]. Here, however, we investigate the mean developmental trajectory numerically which is more efficient than individual-based simulations, especially for the complex developmental trajectories under stage-dependent scenario. In addition, as each organism starts with a single germ-like cell, 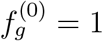 and 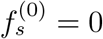. In *ND*^*i*^, cells only produce germ-like cells, thus 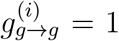 for all *i*, and all other probabilities are irrelevant. Therefore, 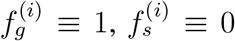. From Eq (8), the growth rate of *ND*^*i*^ is 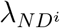 = ln 2. Biologically, it means that organisms double their size per unit of time when they take the *ND*^*i*^ strategy.

## Results

### Stage-dependent cell differentiation promotes irreversible cell differentiation in small organisms

Theoretically, differentiation probabilities at different cell divisions could be arbitrary values. Therefore, differentiation probabilities can be arbitrary values between 0 and 1. However, we rarely observe drastic changes in cell differentiation probabilities during an organism’s development – instead, these probabilities change slowly during development. For instance, cells in a series of closely relative species in *Volvox* family show gradual degrees of germ-soma differentiation Matt and Umen [2016]. Thus, it is natural to assume that the maximum value of the change of two successive differentiation probabilities is small. Therefore, we restrict attention to a small range of *δ* and set *δ* = 0.1 (i.e. 0 ≤ *δ*_*i*_ ≤ *δ* = 0.1, *i* = 1, …, *n*) in this section and the effects of large *δ* will be investigated in the third section. We found that stage-dependent differentiation promotes the evolution of irreversible strategies *ID* compared with stage-independent differentiation *ID*^*i*^ in small organisms, see Fig 2. Specifically, stage-dependent differentiation *ID* evolves at more parameter space of differentiation benefits and costs than stage-independent *ID*^*i*^ in small organisms. However, stage-dependent *ID* gradually loses its advantages when organismal size increases. It has been shown that stage-independent differentiation *ID*^*i*^ is more likely to evolve in large organisms, see *ID*^*i*^ in Fig 2. The conclusion about *ID*^*i*^ is consistent with the previous findings Gao et al. [2021].

**Figure 2:**
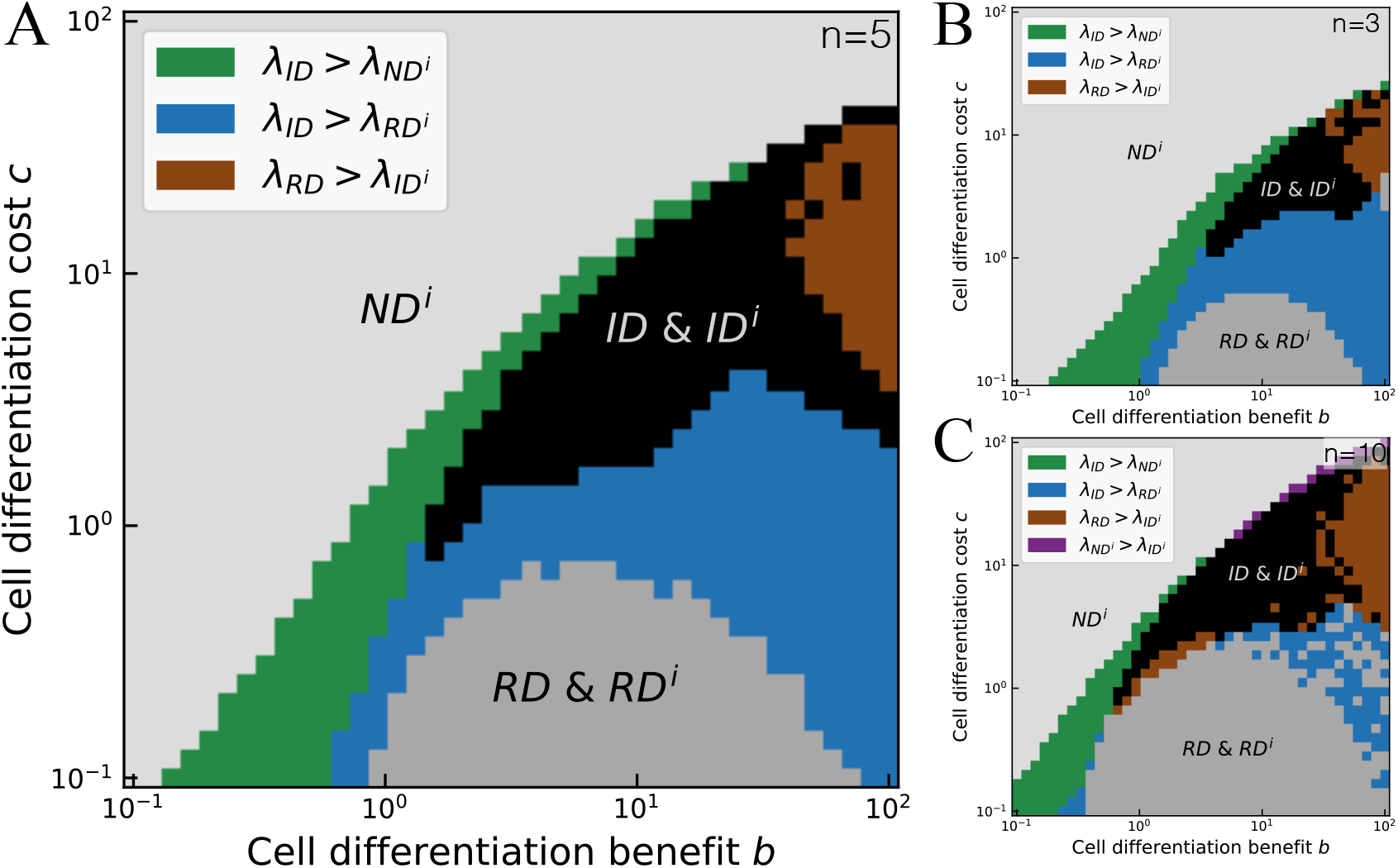
Comparison of optimal strategies between stage-independent and stage-dependent differentiation. Comparison of the parameter space of the optimal strategy between stage-independent and stage-dependent cell differentiation under maximal cell number of division rounds of *n* = 5 (panel **A**), *n* = 3 (panel **B**) and *n* = 10 (panel **C**). The grey, dark grey, and black areas represent the parameter space where the optimal strategies are the same under both stage-independent and stage-dependent cell differentiation. The green strip represents stage-dependent *ID* leading to a larger growth rate than stage-independent *ND*^*i*^. Similarly, the blue area and the brown area represent *ID* and *RD* leading to higher growth rates than stage-independent *RD*^*i*^ and *ID*^*i*^, respectively. Purple color represents *ND*^*i*^ leads to a higher growth rate than *ID*^*i*^ in panel **C**. Parameters of all panels: 0 ≤ *δ*_*i*_ ≤ 0.1, and *α* = *β* = 1. Parameters of calculating optimal strategy: the number of initial sampling *d*^(1)^, *M* = 1000, the number of stage-dependent strategies starting with a given *d*^(1)^, *R* = 100, for more detail, see S2 Appendix. At each pixel, the frequency of each optimal strategy was calculated across 100 replicates in panel **A** and 20 replicates in the rest panels.

Next, with the constraint of *δ*, we investigate the effects of stage-dependent differentiation on an organism’s growth rate under varying differentiation benefits and costs, comparing it with the results of stage-independent differentiation strategies. We first focus on the parameter space where both stage-independent and stage-dependent differentiation evolve the same strategies. *ND*^*i*^ dominates under both stage-independent and stage-dependent differentiation at high differentiation costs which largely decreases an organism’s growth rates under differentiation strategies (*RD*^*i*^, *ID*^*i*^, *RD* and *ID*). In the absence of differentiation benefits, i.e. *b* = 0 and *c* > 0, we show that *ND*^*i*^ is optimal analytically (S4 Appendix). Additionally, if there is only a single cell division (*n* = 1), *ND*^*i*^ is still optimal in the absence of differentiation costs, i.e. *c* = 0 and *b* > 0 (S4 Appendix). This is because differentiation benefits 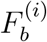 at the *i*th division are based on the fraction of soma-like cells after the (*i* − 1)th division. Similarly, under the scenario of high costs and low benefits, stage-dependent differentiation bears huge costs, thus only *ND*^*i*^ is chosen. When benefits are much higher than costs, then reversible differentiation (*RD*^*i*^, *RD*) is chosen under both stage-independent and stage-dependent differentiation. This is because differentiation benefits will cover the differentiation costs caused by cell differentiation among divisions, see S5 Appendix for the optimal strategy under larger scales of benefits and costs.

Then, we analyze the effects of organism size on the effects of the occurrence of stage-dependent *ID*. In the model, cell differentiation plays a dual role in growth rate. It provides benefits, but also incurs costs on the growth rate. The best strategy is the one that can maximally use cell differentiation benefits and at the same time reduce costs. So under the conditions of high benefits or high costs, only *ND* is selected. Due to the randomness of cell differentiation probabilities, stage-dependent strategies contain the one that can adjust the fraction of germ-like cells to gain differentiation benefits and avoid differentiation costs during growth, especially in large organisms that contain more cell divisions, see Fig 2. In small organisms, due to the constraints on the fluctuations of two successive cell differentiation probabilities, stage-dependent *ID* strategies only accumulate limited differentiation benefits. Since stage-dependent *ID* needs to undergo the differentiation from germ-like to soma-like first and then at least cell type turns irreversible, thus higher cell differentiation costs, especially high differentiation from germ-like to soma-like, will prohibit it from being the optimal strategy. But for *ID* in large organisms that need more cell divisions to mature, the differentiation from germ-like to soma-like can occur only in the first several cell divisions to gain benefits, then cells can remain irreversible to avoid costs in the following cell divisions cells. Thus, stage-dependent differentiation strategies (either *RD* or *ID*) can lead to higher growth rates than stage-independent ones (either *RD*^*i*^ or *ID*^*i*^) in small organisms for the account of their flexible adjustment of the differentiation probability patterns.

### Irreversible germ differentiation dominates among optimal stage-dependent irreversible cell differentiation

To further analyze why stage-dependent irreversible differentiation is favored over stage-independent irreversible differentiation in small organisms, we should further study the possible irreversible differentiation forms in *ID*. In the model, an organism can contain two cell types, thus irreversibility can occur on either cell type. Therefore, the stage-dependent irreversible differentiation (*ID*) can further be classified into three subcategories: irreversible germ differentiation *IGD* (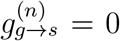and 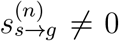), irreversible soma differentiation *ISD* (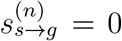and 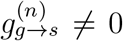), and irreversible germ and soma differentiation 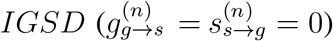. Next, we investigate the occurrence conditions of each sub-strategy.

The results show that among the optimal *ID* in small organisms, *IGD* evolves at most parameter space of benefits and costs, see Fig 3 A and B. *IGD* leads stage-dependent *ID* replaces *ND*^*i*^ as the optimal strategy in small organisms at small *c*, see Fig 2A and Fig 3B. Specifically, we first found that the *IGD* strategy replaces *ND*^*i*^ when *b* is slightly larger than *c*. Under this scenario, the best strategy would be to produce a few soma-like cells to use differentiation benefits, but decrease the differentiation probabilities between cell types to avoid differentiation costs as the growth rate is a tradeoff between differentiation benefits and differentiation costs based on Eq (8). Thus, the *IGD* that produces few soma-like cells in the first few cell divisions and then turns into irreversible becomes optimal (the first panel in Fig 3C). The *IGD* strategy can keep a high fraction of germ-like cells which increases the growth rate by increasing the number of offspring i.e. 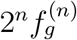. Under this *IGD* strategy, although the differentiation probabilities of soma-like cells 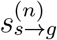 is not small, we should note that the differentiation costs are still low as the number of soma-like cells is small, which is 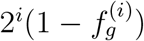 after the *i*th cell division. Then, we found that *IGD* is optimal when *b* is much larger than small *c* in small organisms (the second panel in Fig 3C). Under this scenario, due to the tradeoff between differentiation benefits and costs, the *IGD* strategy with higher germ-like differentiation probabilities *g*_*g*→*s*_ at first several cell divisions becomes optimal. Taken together, we found that *ID*’s sub-strategy *IGD* evolves at low *c*. Furthermore, we show an analytical proof that except for *n* = 1 (S4 Appendix), either *RD* or *ID* is optimal in the absence of cell differentiation costs, i.e. *c* = 0 and *b* > 0 (S6 Appendix). The finding indicates that without the punishment of differentiation costs *ND* cannot be selected.

**Figure 3:**
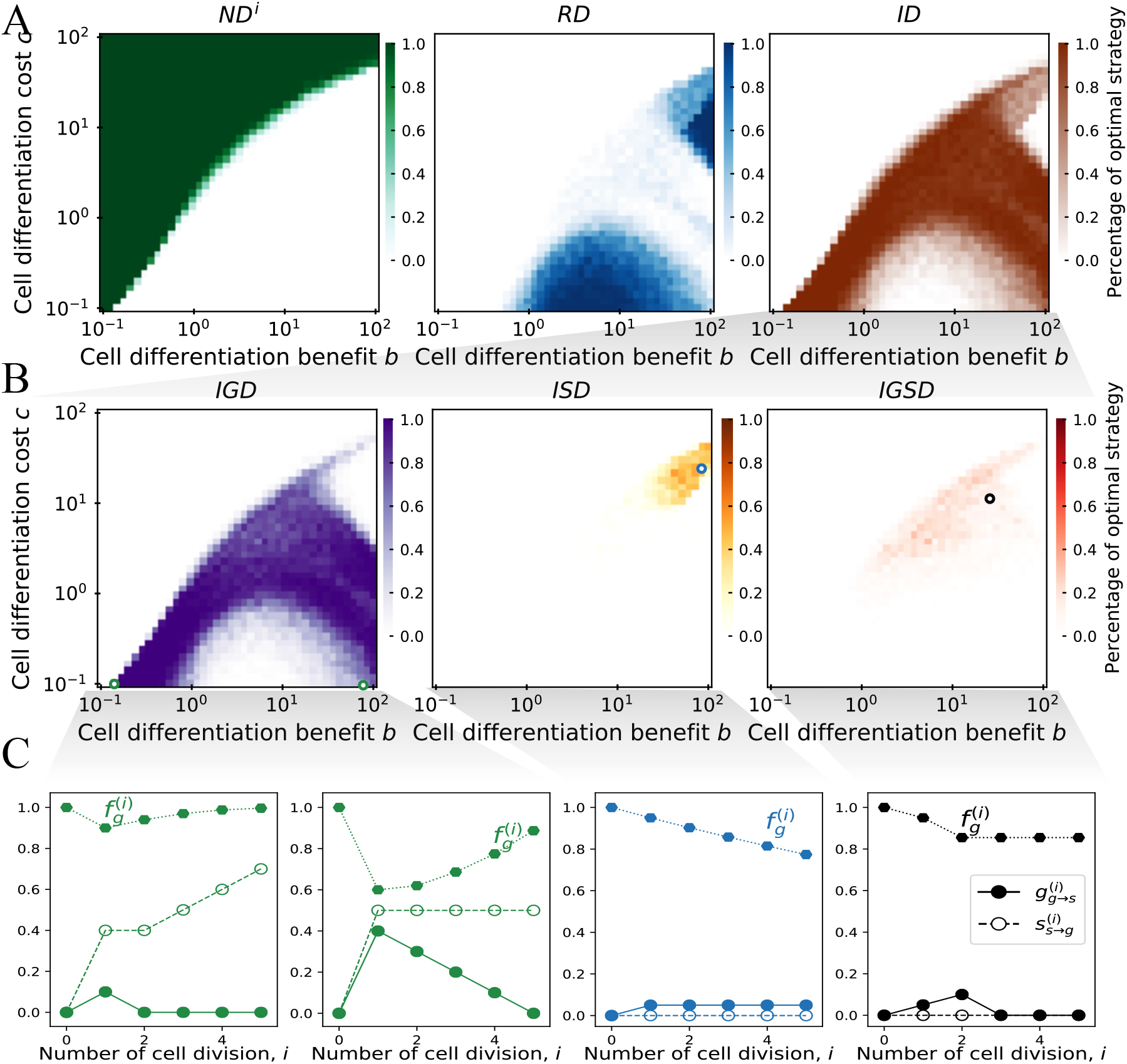
Irreversible germ differentiation evolved mostly among irreversible differentiation under stage-dependent differentiation. **A**. Fractions of three stage-dependent cell differentiation strategies being optimal under differentiation benefits and costs. **B**. Fractions of three sub-irreversible stage-dependent strategies being optimal under differentiation benefits and costs. **C**. Cell differentiation probabilities 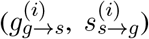 and the frequencies of germ-like cell 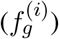 of the optimal irreversible strategy via cell divisions at the parameter space indicated by circles in panel **B**. The circle color follows that in Fig 3 Parameters of all panels: maximal cell number of division rounds *n* = 5, 0 ≤ *δ*_*i*_ ≤ 0.1, and *α* = *β* = 1. At each pixel, the frequency of each optimal strategy was calculated across 100 replicates. Parameters of calculating optimal strategy: the number of initial sampling *d*^(1)^, *M* = 1000, the number of stage-dependent strategies starting with a given *d*^(1)^, *R* = 100, replicates for each pixel is 100, for more detail, see S2 Appendix.

Meanwhile, We found that *ISD* and *IGSD*, the other subcategories of *ID*, evolve at both intermediate values of differentiation benefits and costs, see the last two panels in Fig 3B. We analytically proved that both cell differentiation benefits and costs are indispensable factors for the evolution of *ISD* and *IGSD*, see the proof in S7 Appendix. The subcategory strategy *IGSD* and *ISD* of *ID* evolves at both high *b* and high *c*. Specifically, *IGSD* evolves at both higher *b* and *c* than the strategy of *ISD*. This is an account of the differences in irreversibility features of cell types between *IGSD* and *ISD*. Compared with *ISD* strategies, *IGSD* with both irreversible cell types at last cell division bears lower cell differentiation costs, thus it can evolve either at higher *c* or at low conditions of *b* and *c* than *ISD* (the last two panels of Fig 3C). Meanwhile, *IGSD* has relatively higher fractions of germ-like cells than *ISD*, which leading a larger number of offspring i.e. 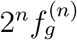 and then leads to a higher growth rate based on Eq (8). Additionally, it is noteworthy that for the evolution conditions of stage-independent irreversible soma differentiation *ISD*^*i*^, the only *ID*^*i*^ under stage-independent cell differentiation, the result is consistent with our previous study Gao et al. [2021].

In addition, compared with the previous work which investigated cell differentiation under stage-independent differentiation Gao et al. [2021], stage-dependent differentiation also promotes the evolution of irreversible differentiation under the effects of *α* and *β*, see S8 Appendix. Meanwhile, we found that under stage-dependent differentiation *α* plays a similar role as that of stage-independent differentiation. That is, irreversible differentiation strategies are optimal when *α* < 1, i.e. the cell division rate component *F*_*b*_ accelerates with *α*, see Fig 8 in S8 Appendix. However, the effects of *β*, measuring the relative weight of cell transition between germ-like and soma-like cells, leads to different results between stage-dependent and stage-independent differentiation. Specifically, we found that irreversible differentiation evolves across all values of *β*. As the subcategory *IGD* of *ID* evolves when *β* is small, and *ISD* and *IGSD* of *ID* evolve when *β* is large.

### Large changes in two successive probabilities of cell differentiation prevent irreversible differentiation from becoming optimal

Without constraints, the differences of cell differentiation probabilities in successive cell divisions can take any value, i.e. 0 ≤ *δ* ≤ 1, where 0 ≤ *δ*_*i*_ ≤ *δ* ≤ 1. Then, cells’ differentiation probabilities and related organism’s growth are completely arbitrary. An extreme example that optimally exploits the potential of somatic cells would be that both types of cells produce soma-like cells in the first (*n* − 1) divisions, and then all produce germ-like cells in the last division. We refer to this cell differentiation as “extreme differentiation” (*ED*). We will first take *ED* as a typical example to investigate the effects of stage-dependent differentiation without any constraints. *ED* is a strategy that fully uses cell differentiation benefits. In contrast, *ND*^*i*^ is a strategy that does not receive cell differentiation benefit or cell differentiation cost. Next, we compare the evolving conditions for *ED* and *ND*^*i*^. Naturally, we expect that when *b* ≫ 1 and *c* ≪ 1, *ED* is optimal, and when *c* ≫ 1 and *b* ≪ 1, *ND*^*i*^ is optimal. Based on Eq (8), the growth rate of *ED* is 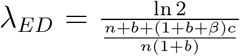 and the growth rate of *ND* is 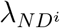 = ln 2. Thus, when 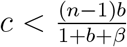, we have *λ*_*ED*_ > 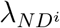 and when 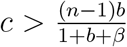 we have *λ*_*ED*_ < 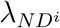, see Fig 4. The outcome indicates that if the costs of differentiation are high, the strategy of no differentiation will be chosen, and if the benefits of differentiation are high, differentiation strategies will be selected.

**Figure 4:**
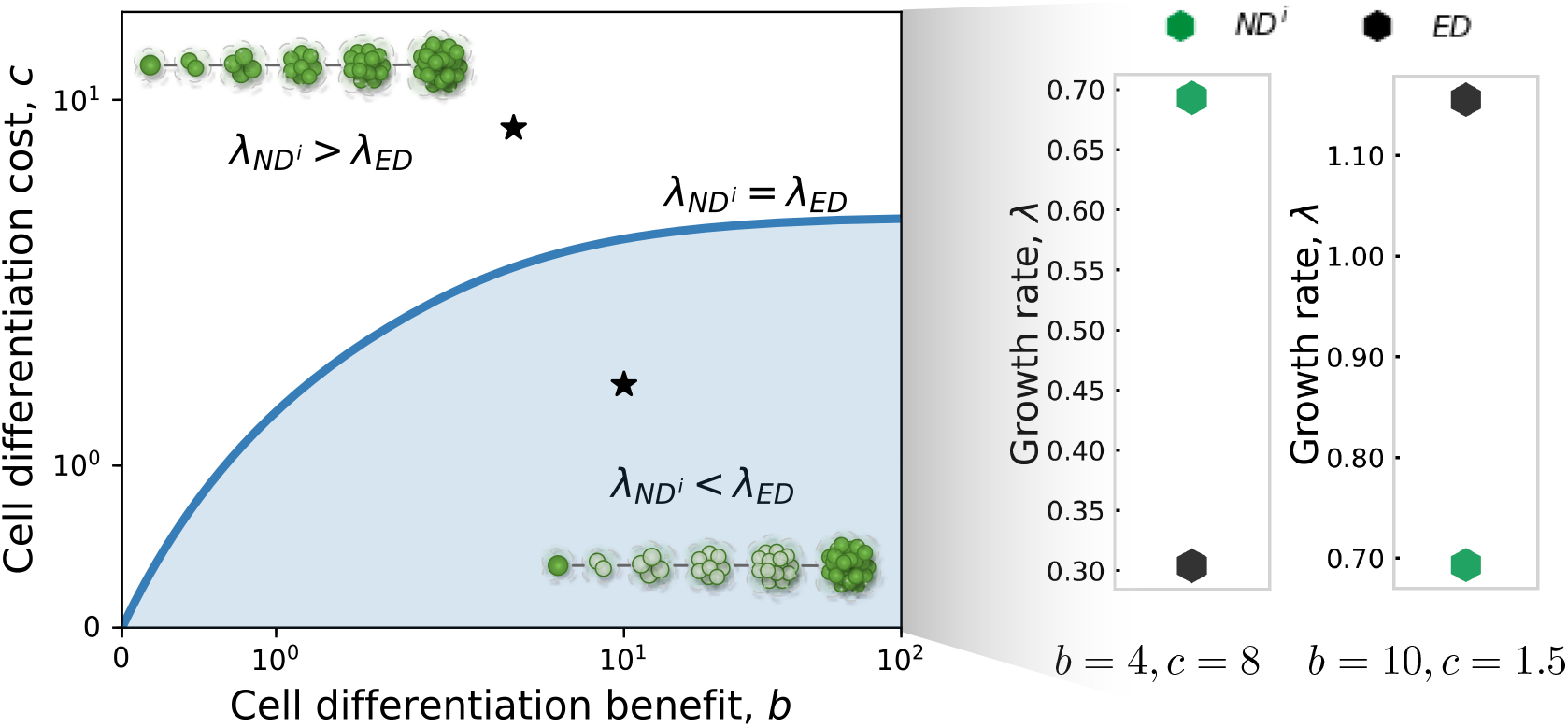
The evolutionary conditions for non-differentiation *ND*^*i*^ and extreme differentiation *ED*, and their corresponding growth rates. The blue line represents the condition for *λND*^*i*^ = *λ*_*ED*_. The shaded area represents where *λND*^*i*^ < *λ*_*ED*_. We found that *ND*^*i*^ is optimal under high *c* and *ED* is optimal under high *b*. The black stars correspond to the parameter combinations where growth rates have been calculated in the right panel. Parameters: *n* = 5 and *α* = *β* = 1.

We next investigate the effects of the maximum change of two successive differentiation probabilities i.e. parameter *δ* on the growth rates of the general stage-dependent strategies *ND*^*i*^, *RD*, and *ID*. We found that irreversible differentiation cannot be optimal under large *δ*, see Fig 5. Large *δ* means more randomness of cell differentiation during an organism’s growth. A higher value of *δ* intensifies the spectrum of an organism’s growth rate except for *ND*^*i*^ whose differentiation probabilities don’t change with *δ*. For instance, we found that the growth rate of the optimal differentiation strategies including *RD* and *ID* all increase with increasing *δ*. However, *RD* has a relatively greater increase than *ID* (Fig 5). Furthermore, we found that *RD* outcompetes *ID* and turns into the optimal strategy when *δ* is 1. Specifically, when *δ* = 0.1, we found that the sub-strategy *IGD* of *ID* leads to a larger growth rate than *RD*, whereas when *δ* = 1, *RD* outcompetes *IGD* and leads to the largest growth rate, see Fig 5 A-C and D-F. Based on Eq (8), we know that the growth rate depends both on cell division rates and the number of offspring (the fraction of germ-like cells after the *n*th division). Furthermore, the cell division rate is proportional to the fraction of soma-like cells but inversely proportional to the differentiation probabilities which cause differentiation costs. Taken together, the largest growth rate favors the strategy with a higher fraction of soma-like cells all the time, a higher fraction of germ-like cells after the last cell division, and lower differentiation probabilities. *RD* contains the strategy to increase the fraction of soma-like cells in the middle stages of cell divisions and the number of offspring which is the number of germ-like cells after the *n*th cell division. Thus, higher *δ* prohibits the emergence of strategy *ID* being optimal.

**Figure 5:**
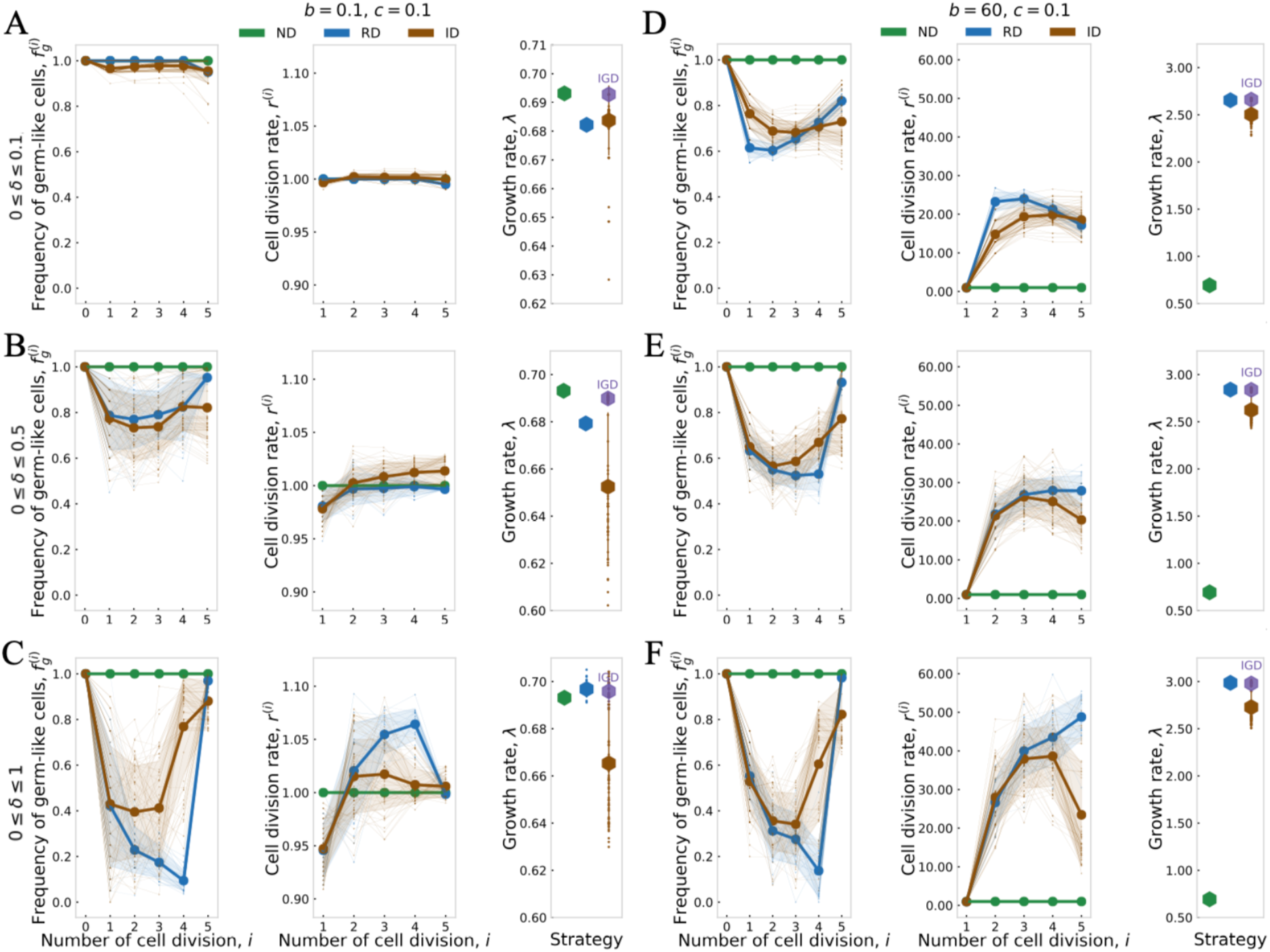
The effect of the maximum change of two successive differentiation probabilities *δ* on the growth rate of optimal strategies. Frequencies of germ-like cells, cell division rates, and growth rate of the optimal stage-dependent strategy of each category under *δ* = 0.1, *δ* = 0.5 and *δ* = 1 respectively, *i* = 1, 2, …, *n*. Small dots are the values of each interesting feature at each cell division. Thick lines are the averaged values at each cell division. The shaded areas indicate the standard deviation. *α* = *β* = 1 and colors correspond with those in Fig 3. Parameters: *n* = 5, and *α* = *β* = 1, we chose 20 duplicates for generating the optimal strategies in each category which includes subcategories.

## Conclusion and discussion

We investigated the effect of stage-dependent differentiation on an organism’s growth and compared it with stage-independent cell differentiation. Stage-independent cell differentiation only allows a fixed cell differentiation probability for a cell type. Stage-dependent differentiation, by contrast, refers to being capable of changing differentiation probabilities in consecutive cell divisions. The most extreme case would be an organism that is entirely consisting of soma-like cells until the last cell division, where all cells turn into germ-like cells to produce as many offspring as possible. Stage-dependent differentiation intensifies the fluctuation of the germ-soma ratio during an organism’s growth, which further increases the complexity of competition between different strategies. We used the growth rate of an organism as a proxy to investigate the growth competition of different strategies under different benefits and costs. Based on the differentiation probabilities in the last division, we classify stage-dependent differentiation into three categories: non-differentiation *ND*^*i*^, reversible differentiation *RD*, and irreversible differentiation *ID*. The evolution of irreversible differentiation under stage-independent differentiation has been demonstrated by previous work to be challenging Gao et al. [2021]. Contrary to our expectations, we found that stage-dependent differentiation favors *ID* (in the last division step) more than stage-independent irreversible differentiation *ID*^*i*^ in smaller organisms. Specifically, *IGD*, a sub-strategy of *ID*, leads to a higher growth rate than other strategies in small organisms. Additionally, *ISD* and *IGSD* evolved in the parameter space with intermediate benefits and costs, consistent with previous findings Gao et al. [2021]. Finally, we found that large differentiation probability variation prohibits irreversible differentiation *ID* from becoming the optimal strategy. The findings indicate that stage-dependent differentiation favors the evolution of irreversible differentiation in small organisms and with limited variations between successive cell divisions.

That irreversible differentiation is favored in small organisms is contrary to the intuition provided by stage-independent differentiation, where irreversible differentiation is favored in large organisms Gao et al. [2021]. Our previous work has shown that the minimum size for irreversible differentiation occurring is *n* = 6 Gao et al. [2021]. This discrepancy arises because of the flexibility of the developmental trajectories under stage-dependent differentiation. These complex developmental trajectories in different categories increase the growth differences between different strategies. Thus, we found that the optimal strategies of differentiation categories can lead to divergent growth rates. In addition, stage-dependent irreversible differentiation evolves two more subcategories than stage-independent one: irreversible germ differentiation *IGD* and irreversible germ and soma differentiation *IGSD*. The broad form of stage-dependent differentiation strategies can capture more cell differentiation patterns in reality. For example, the evolution of *IGSD* can help us to understand cell lineage segregation in nature Matt and Umen [2016]. Our model can screen the stage where irreversible differentiation emerges, in line with the question of early segregation of germ and soma in animals Buss [1983], Knaut et al. [2000], Buehr [1997], McLaren [2003], Extavour and Akam [2003], but late in most plants Lanfear [2018]. To identify the segregation, we need to investigate the irreversible developmental states of germ-like and soma-like cells in our model. Future work is necessary for seeking and analyzing the conditions where different segregation occurs.

Previous investigations of cell differentiation mostly focused on the state with a group of undifferentiated clonal cells Michod [2007], Gavrilets [2010], Rodrigues et al. [2012], Goldsby et al. [2014], Cooper and West [2018], Yanni et al. [2020], Liu et al. [2021], Cooper et al. [2021, 2022] or cells with randomly chosen initial cell types (similar to aggregated organisms) Rodrigues et al. [2012]. The focus of these studies was on the final static conditions that lead to the division of labor rather than the dynamic process during an organism’s growth. These models ignored the dynamic developmental trajectories of organisms from newborn to maturity. In our model, the developmental trajectories of each organism are recorded by stage-dependent differentiation probabilities, allowing us to know the dynamic fractions of each cell type during an organism’s growth, which further allow us to investigate cell differentiation patterns. In addition, Rodrigues et al. have considered cell differentiation probability as an evolving trait to understand the evolution of differentiation Rodrigues et al. [2012]. They concluded that differentiation costs, compared with the difference in division rates between cell types, have less impact on the evolution of terminal and reversible differentiation. They also found that differentiation costs played a crucial role in the evolution of diversity differentiation strategies. Moreover, Rodrigues et al. investigated developmental strategies in filament multicellular organisms with two essential tasks, and they found that high differentiation costs can promote the evolution of symbioses. In the model, we employ functions to demonstrate differentiation benefits and costs (Eq (3), Eq (4) as they can capture more general forms of benefits and costs by varying relevant parameters.

In our model, since we focus on the evolution process that cells reach the final specialized types, thus we assumed that differentiation occurs randomly and both cell types are capable of cell differentiation Gao et al. [2021]. The assumption is based on the cell differentiation situation of species observed in genus *Volvox*, which reveals that cell types undergo an intermediate and partial differentiation stage in some closed related species before eventually becoming specialized cell types Matt and Umen [2016]. We classify the stage-dependent differentiation strategy based on its differentiation probability at the last round of cell division. The classification is based on the idea that the differentiation strategy (reversible and irreversible) describes the changes in differentiation capability along the cell division process. Nevertheless, we stress that this classification is imperfect, especially for large organisms with more cell divisions, where a more refined classification criterion is needed. However, owing to the simple classification, the current classification can still largely reflect the evolving situation of the specific strategies interested. For instance, the strategy that cells all turn into specialized types after half a round of cell divisions is a subset strategy of *ID*, thus it can only evolve in the parameter space that *ID* emerged. Meanwhile, we assumed that organisms are clonal, growing from a single founding cell. The reasons for our clonal assumption are that multicellularity is formed commonly by clonal division rather than cell aggregation Fisher et al. [2013], Grosberg and Strathmann [1998], Tarnita et al. [2013], Brunet and King [2017], Pentz et al. [2020], Márquez-Zacarías et al. [2021a]. and clonal organisms with identical genes have advantages at purging deleterious mutations and reducing conflicts among cells Grosberg and Strathmann [1998, 2007]. Therefore, clonal multicellularity is predicted to be evolutionarily stable Mikhailov et al. [2009]. In the cell differentiation models of aggregated multicellularity, a relatedness parameter can be used to evaluate the level of cooperation between cell types Ispolatov et al. [2012], Cooper and West [2018], Madgwick et al. [2018], Liu et al. [2021]. Additionally, the maturity size is fixed in the model as previous work has shown that selection favors life cycles where all organisms grow to the same size and fragment into pieces with the same pattern Pichugin et al. [2017]. The assumption is generally in line with the size observation in some species such as *Volvox* Matt and Umen [2016].

We assumed that cell differentiation costs influence an organism’s growth. In nature, cell differentiation and cell plasticity usually originally occur under severe environmental conditions, indicating a differentiation cost involved Gallon [1992], Claessen et al. [2014], Aguirre et al. [2005], Loenarz et al. [2011]. Differentiation cost has been considered in previous theoretical research via varying forms DeWitt et al. [1998], Gavrilets [2010], Ispolatov et al. [2012], Rodrigues et al. [2012], Goldsby et al. [2012], Staps and Tarnita [2022]. But the modeling purpose of cell differentiation costs is the same, i.e. reducing an organism’s fitness. In our model, we are interested in the relative growth advantage between different differentiation strategies. Therefore, we assume that differentiation costs affect the growth rate, reducing cell division rates. Finally, we suppose that cells undergo synchronous cell divisions. This is not true for large multicellularity with many more cell divisions Newport and Kirschner [1982], Matt and Umen [2016]. Asynchronous cell division has been explored under stage-independent differentiation in previous studies, leading to the same predictions as the synchronous one Gao et al. [2021]. Yet, it still needs to be investigated whether asynchronous cell division leads to the same conclusion as synchronous ones under stage-dependent differentiation in the future. Our model could be further extended by including cell death or differentiation costs related to the risk of organism death. Yet, our model gives first insights into understanding the effects of dynamic differentiation on the evolution of cell differentiation in multicellularity.

## S1 Appendix. Growth rate

In our model, we treat both the number of cells and growth time as continuous, distilling the stochastic process down to two quantities for calculating the growth rate: the expected offspring number of germ-like cells *N* and the amount of growth time for an organism to grow *t*. The expected growth rate *λ* can be calculated by the following equation approximately

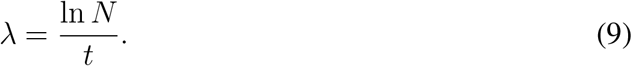

The robustness of the approximation is tested in S3 Appendix. Here, we use *f*_*g*_(*i*) and *f*_*s*_(*i*) to denote the fractions of germ-like cells and soma-like cells after the *i*th cell division. Since each organism starts with a single germ-like cell, *f*_g_ (0) = 1 and *f*_s_ (0) = 0. We use 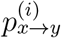 to denote the transition probability from cell type *x* to *y* in the *i*th cell division, where *x* and *y* are either germ-like cells or soma-like cells. Based on Eq (1), we have

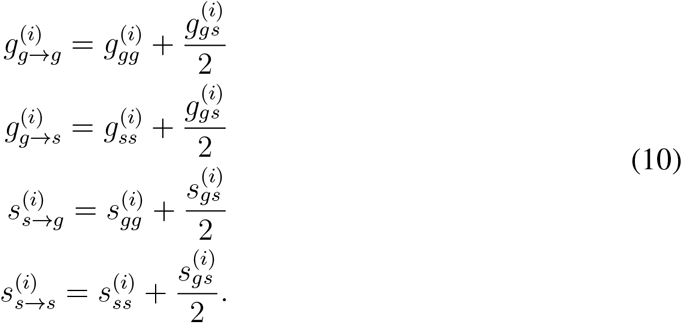

After the *i*th cell division, the expected fraction of germ-like cells is 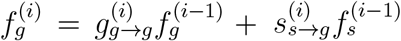 and the expected fraction for soma-like cells is 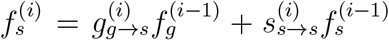 which can be expressed in

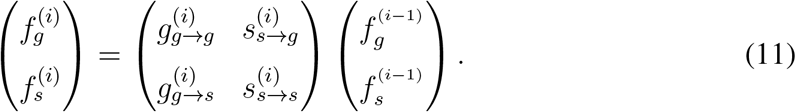

The expected 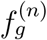 and 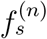 can be calculated recursively by Eq (11)

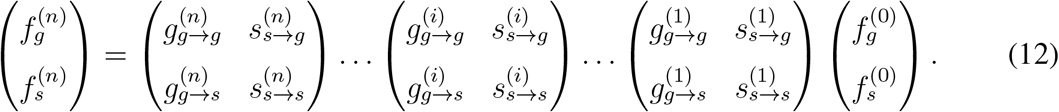

Since cells divide synchronously and no cell dies during growth, the expected number of germ-like cells 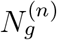 and soma-like cells 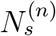 after the *n*th cell division are

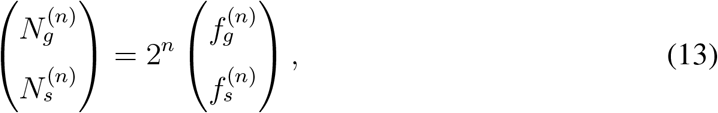

where 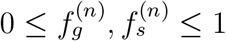.

The cell division rate determines the growth duration of organisms. Since cells divide with a rate 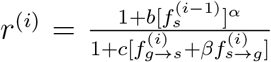 during the *i*th cell division, the waiting time for a cell division *t*^(*i*)^ follows the exponential distribution 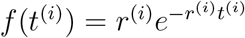, where 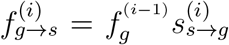 *s*^(*i*)^ and 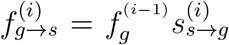,see Eq (6). Thus the expected waiting time from the *i*th cell division to the (*i* + 1)th cell division is 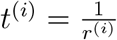 The expected growth time for organisms with total *n* cell divisions is

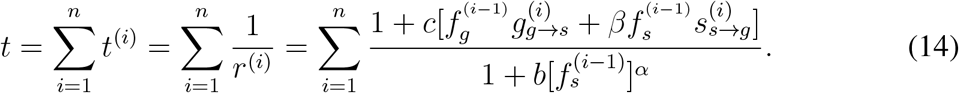

Substituting Eq (13) and Eq (14) into Eq (9), we have

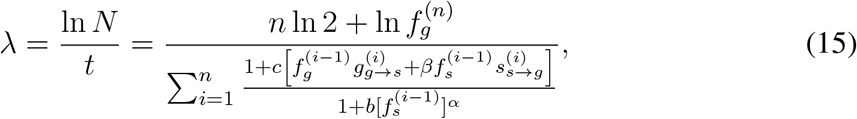

where *n* is the number of total cell divisions of organisms, 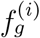 and 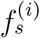 are fractions of germ-like cell and soma-like cell after the *i*th cell division, 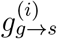 and 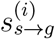 are the transition probabilities between germ-like cell and soma-like cell at the *i*th cell division (1 ≤ *i* ≤ *n*). We have 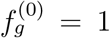 and 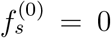. For the non-differentiation strategy *ND*^*i*^, no soma-like cells are produced during growth, i.e. *g*_*g*→*g*_ = 1 and *g*_*g*→*s*_ = *s*_*s*→*g*_ = *s*_*s*→*s*_ = 0. Therefore, 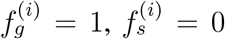. Thus from Eq (15) the growth rate of *ND* which is denoted by 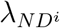 is ln 2. Biologically, the growth rate of *ND*^*i*^ describes the number of cells doubling per unit of time. As we defined strategies based on the series of cell differentiation probabilities, the growth rate of a strategy should be a distribution rather than a fixed value. But for calculation convenience, we took Eq (15) as an approximation of a stochastic differentiation strategy. In appendix S3, we show the approximation is reliable in finding the optimal differentiation strategy.

## S2 Appendix. Numerical calculation of the growth rate of the stage-dependent differentiation strategy

We introduce the method of calculating growth rate numerically in stage-dependent cell differentiation. To find the optimal strategy, at a fixed benefit and cost condition, we use the Monte-Carlo methods randomly to sample differentiation strategies and then calculate and compare the growth rates of organisms under these strategies. Grid search is used to find the optimal strategy in different conditions of benefits and costs. We first look at the cell differentiation probabilities in the first division step 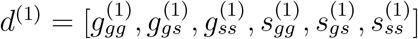. Each probability can take the value 0 or other values by increasing 0.1 from 0 at each time until reaching the highest value 1, thus there are 11 possible values for each probability, i.e 0, 0.1, 0.2, …, 1. We first define the number of probability combinations. Since 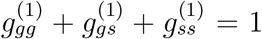 and 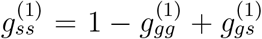, as long as we know the values of 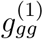 and 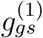, we know *g*_*ss*_. When *g*_*gg*_ = 0, *g*_*gs*_ can take the 11 values from 0 to 1, Thus, there are totally 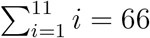 combinations for 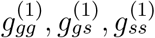 The same number of combinations exist for soma-like cells. Thus, there are a total of 66 *×* 66 = 4356 combinations for *d*^(1)^. As long as *d*^(1)^ is chosen, we need to identify *d*^(2)^, *d*^(3)^, …, *d*^(*n*)^. *d*^(2)^ deviates from *d*^(1)^ by either a 0 or *δ*_2_. That is 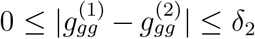. The same for the other probabilities *g*_*gs*_, *g*_*ss*_, *s*_*gg*_, *s*_*gs*_, *s*_*ss*_. The choice of *d*^(*i*+1)^ depends on number of neighbours of *d*^(*i*)^, which further depends on the elements *d*^(*i*)^. For *δ*^(*i*+1)^ = 0.1, if *g*_*gg*_ = *s*_*ss*_ = 1 in *d*^(*i*)^, then *g*_*gg*_ and *s*_*ss*_ can only be decreased or be constant. Thus, there are 5 choices for *d*^(*i*+1)^. However, if the elements in *d*^(*i*)^ are either 0.3 or 0.4, then each element can be increased, decreased, or unchanged. Therefore, it has 13 choices for choosing *d*^(*i*+1)^. Let’s take *d*_1_ = [0.3, 0.3, 0.4, 0.3, 0.3, 0.4] as an example, then it’s neighbours are

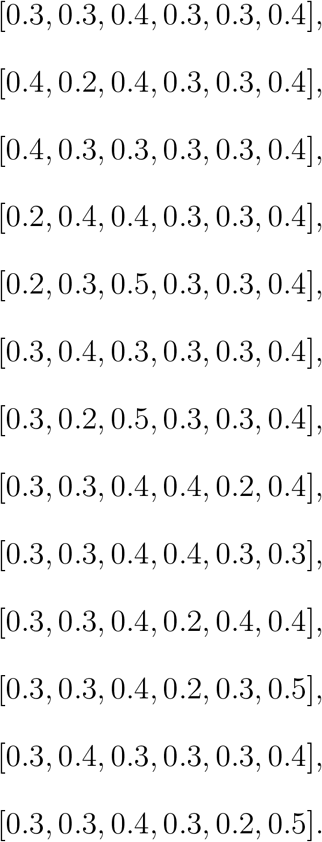

Note that *d*^(1)^ is considered as one of its neighbours. To generate a stage-dependent differentiation strategy, we first chose *d*^(1)^ from the combination pool and then chose *d*^(2)^ from *d*^(1)^’s neighbors and repeat the process until obtaining *d*^(*n*)^. We choose each strategy randomly following a uniform distribution. As long as we have classified strategies, we will have a pool of each strategy and then we choose strategies from the pools. Specificity, we first choose the last probabilities and then randomly choose other probabilities backward in rounds of cell division. For example, for choosing a *RD* strategy, we first randomly pick the probabilities at the *n*th round of cell division, which should satisfy 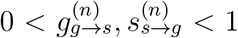 Then we randomly choose the probabilities at the (*n* − 1)th round of cell division and so on until the first one.

In stage-independent cell differentiation, we calculate the growth rates of each strategy in the cell differentiation probabilities pool. We seek the optimal strategy which leads to the fastest growing among these 4356 strategies. To find the optimal strategy at a given parameter point, we first chose *M* = 1000 values for *d*^(1)^ from the cell differentiation pool. Then for each chosen *d*^(1)^, we randomly chose *R* = 100 stage-dependent strategies, all generated from this *d*^(1)^. *R* = 100 is the sampling size of the stage-dependent strategies from the same initial differentiation probabilities *d*^(1)^. Then we compute the growth rate of the 10^5^ strategies and choose the strategy leading to the largest growth rate. Next, we optimize that strategy further. For the optimal strategy with the largest growth rate, we compare its growth rate with a slightly modified strategy. The modified strategies include the one removing *d*^(1)^ but compensating with an *d*^(*n*+1)^ or removing *d*^(*n*)^ by compensating with an *d*^(0)^. Specifically, for the focused *D* = [*d*^(1)^, *d*^(2)^, …, *d*^(*n*)^], we check whether *D*^*t*^ = [*d*^(0)^, *d*^(1)^, *d*^(2)^, …, *d*^(*n*−1)^] or *D*^*t*^ = [*d*^(2)^, …, *d*^(*n*)^, *d*^(*n*+1)^] leads to a higher growth rate over *D*. Here the *d*^(0)^ is one neighbour of *d*^(1)^, and *d*^(*n*+1)^ is one neighbour of *d*^(*n*)^. If the *D*^*t*^ leads to a higher growth rate, we keep the process until we find the *D*^*t*^ which makes the growth rate stay at the maxima. We aim to find a local optimum close to the strategy that was identified in our grid search. Local optimization stops when the largest steady growth rate in the local neighbourhood is identified. Overall, we first search the optimal *D* globally by randomly choosing *d*^(1)^, represented by *M* and *R*. The values of *M* and *R* and the number of duplications used in the main text were chosen to ensure the optimal strategy converging to a unique strategy. Then, we used a local grid search by modifying *d*^(1)^ or *d*^(*n*)^ of a strategy until finding the optimal *D*. Besides, we have constructed initial sampling strategies from the middle of *d*^(*i*)^ sequences. We first identified 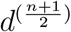 if *n* is odd and 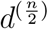 if *n* is even, and then constructed the rest *d*^(*i*)^s. The results show that there is almost no differences in terms of searching for optimal strategies between the two methods.

## S3 Appendix. Robustness of the growth rates of stochastic differentiation strategies

In our model, we calculated the growth rate of a stochastic strategy based on its expected growth time and expected number of germ-like cells. That is, we treat a stochastic differentiation strategy that may contain many potential developmental trajectories as a deterministic one. Theoretically, the growth rate of a stochastic strategy should be a random variable.

Next, we show that the method used in the model is a good approximation for seeking the optimal strategy in an average sense. To simulate the consecutive stochastic differentiation probabilities, at a given stage we need to know the differentiation probability distribution that the next consecutive probabilities follow. Without loss of generality, we assume that the differentiation probabilities follow Gaussian distribution. Then, the coming cell differentiation probability of a cell type is a variable with the last past differentiation probability as the mean. For an arbitrary strategy *D* = [*d*^(1)^, *d*^(2)^, …, *d*^(*n*)^], we can get 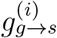 and 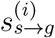 for each *i, i* = 0, 1, …, *n*. Then the variable 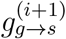 follows the Gaussian distribution 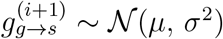, where 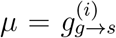.To capture the growth rate of the stochastic differentiation strategy *D*, we randomly choose the new 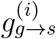 and the new 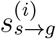 from the Gaussian distribution with mean 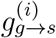 and 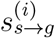respectively, *i* = 1, 2, 3, …, *n*. Each sampling will generate a new strategy *D*^∗^, which is a potential developmental strategy based on *D*. For each *D*^∗^, we can calculate its growth rate based on Eq (15). Then we adopt the Monte Carlo method to capture the potential growth rate distribution by randomly choosing a large number of *D*^∗^ and calculating their growth rate. Based on our numerical calculation, we found that our approximation is along well with the expected growth rate of a randomly chosen strategy (Fig 6A). The value of variance *σ* is undefined. As here we focus on the mean behavior of a strategy, thus variance only impacts the range of growth rate. Furthermore, we testified whether the conclusion under the approximation is consistent with the statistical results introduced above. We found that the optimal strategies are the same (Fig 3 and Fig 6B), indicating the robustness of the approximation method. However, we should note the expected growth rate of a stochastic differentiation strategy may not be equal to our approximation. The former is Σ *λ* _*k*_*p*_*k*_, where *k* is the all possible trajectories of *D*^∗^, *p*_*k*_ is the corresponding probability of choosing trajectory *k*, and *λ*_*k*_ is the growth rate under trajectory *k. p*_*k*_ is multiplication of 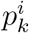 which is the probability of choosing a differentiation probability for either germ-like cell 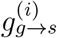 or soma-like cell 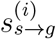 in *D*^∗^, where *i* = 1, 2, …, *n*. In the numerical calculation (Fig 6), we roughly classify 8 intervals i.e. 8 different probabilities for generating 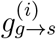 or 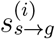 for a given *i*. The 8 intervals are classified based on boundaries of *µ* + *j* ∗ *σ*, where *j* = −3, −2, −1, 1, 2, 3. As we seek the optimal strategy, which depends on the relative difference between different strategies i.e. the rank of the growth rate of different strategies, we employ the approximation to seek the optimal strategy in the model.

**Figure 6:**
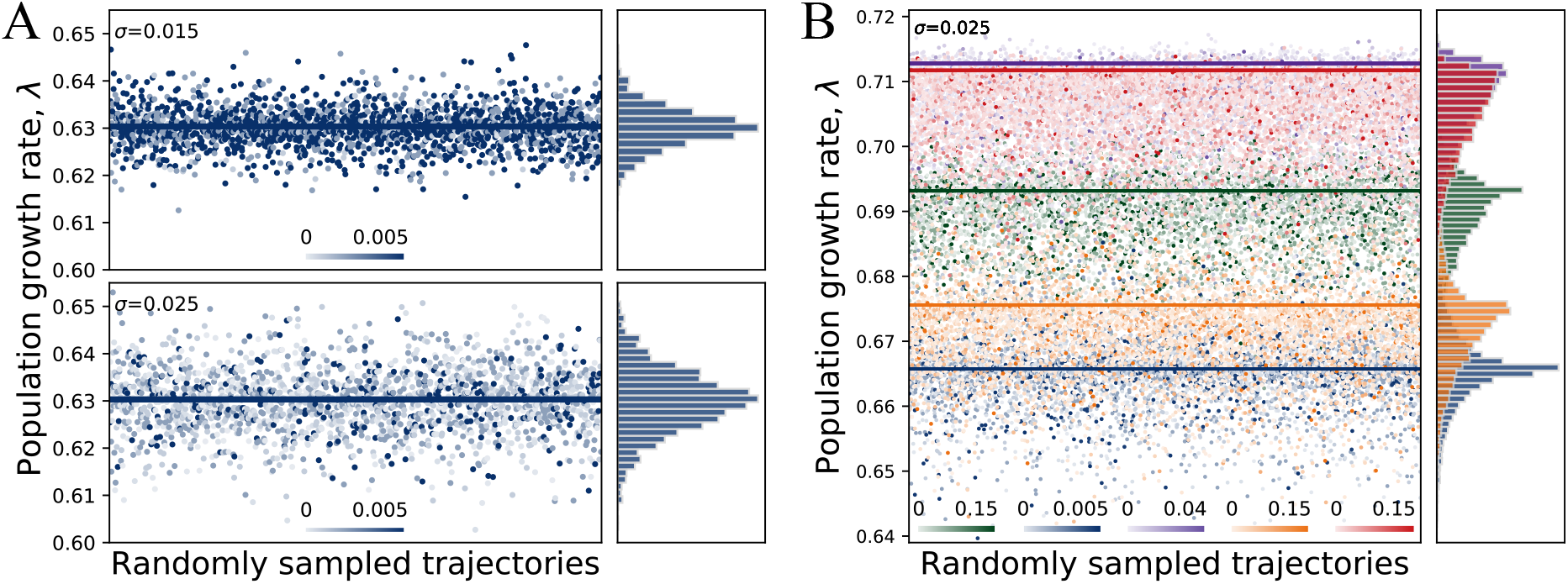
Comparison of growth rates by approximation and random sampling. **A**. Growth rate comparison of potential random trajectories (strategies) of a randomly chosen *RD* strategy under Gaussian distribution with variance 0.015 and 0.025 respectively. The small blue dots represent the potential trajectories. The lines represent the expected growth rates calculated based on Eq (15). The color of the dots represents the probability of the randomly chosen *RD* choosing the dots. The histograms represent the distribution frequency of *λ*. **B**. Growth rate of randomly sampled optimal strategies of each category (*ND*^*i*^, *RD, IGD, ISD* and *IGSD*). The optimal strategy is obtained based on the calculation in S1 Appendix and the grid search method in S2 Appendix. The color of the dots represents the probability of the optimal given strategy choosing the strategy. The histograms represent the distribution frequency of *λ*. The colors represent the same strategy as that in Fig 3. Parameters for all panels *δ* = 0.05, *n* = 5 and *b* = *c* = 1. For calculating the growth rate of each strategy, see the appendix S2 Appendix.

## S4 Appendix. Optimality of non-differentiation strategy *ND*^*i*^

### *ND*^*i*^ is optimal in the absence of cell differentiation benefits for any maximal cell division number *n*

When the cell differentiation benefit is absent, i.e. *b* = 0 and *c* > 0, we find that *ND*^*i*^ is the optimal strategy based on Eq (15).

### *ND*^*i*^ is optimal in the absence of costs for *n* = 1

Next, when there is only one cell division (*n* = 1), we prove that *ND*^*i*^ leads to the largest growth rate under *b* > 0 and *c* = 0. Since 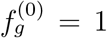 and 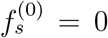, based on Eq (14), the growth time is

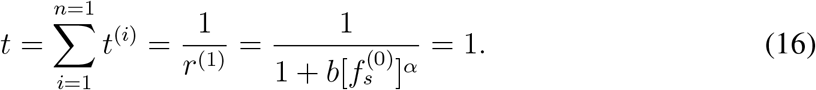

Substituting Eq (16) into Eq (15) and using Eq (12), we have

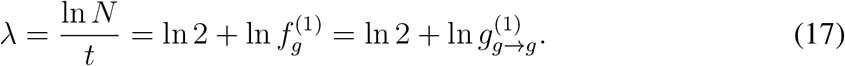

Since 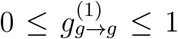, the optimal strategy is *ND*^*i*^ which has 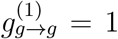. Thus, *ND*^*i*^ is the optimal strategy under *b* > 0, *c* = 0 and *n* = 1.

## S5 Appendix. Optimal strategies of *n* = 5 under a larger range of parameter space

Here, we show that *RD* is optimal when benefits are far larger than costs, see Fig 7. *ND*^*i*^ is optimal when differentiation costs are far larger than benefits.

**Figure 7:**
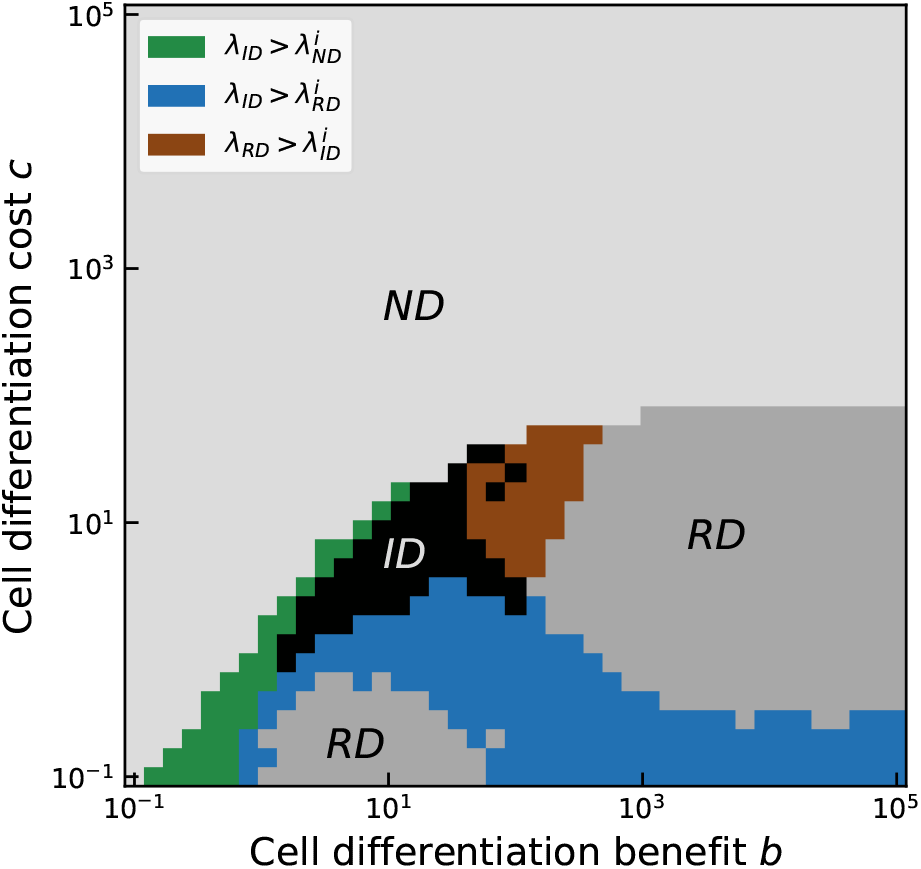
Comparison of the optimal strategies between stage-independent differentiation and stage-dependent differentiation under the large scale of benefits and costs. The colors show the same meaning as that in Fig 2. Parameters: 0 ≤ *δ*_*i*_ ≤ 0.1, *α* = *β* = 1, and *n* = 5. Parameters of calculating optimal strategy: the number of initial sampling *d*^(1)^, *M* = 1000, the number of stage-dependent strategies starting with a given *d*^(1)^, *R* = 100, for more detail, see S2 Appendix. At each pixel, the frequency of each optimal strategy was calculated across 20 replicates..

## S6 Appendix. Either *IGD* or *RD* is optimal in the absence of cell differentiation costs when maximal cell division

*n* > 1. To show the optimal strategy is either *IGD* or *RD* under *b* > 0 and *c* = 0. We first prove *λ*_*IGD*_ > *λ*_*IGSD*_ > 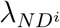, and then prove *λ*_*RD*_ > *λ*_*ISD*_ > 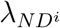.

### *IGD* is optimal among *IGD, IGSD* and *ND*^*i*^

To prove *λ*_*IGD*_ > *λ*_*IGSD*_ > 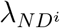, we begin with the proof of *λ*_*IGSD*_ > 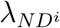. Unlike the *ND*^*i*^ strategy, *ISD, RD, IGD* and *IGSD* are categories which include many strategies. As long as one strategy in *IGSD* has a greater growth rate than *ND*^*i*^, we say *IGSD* is more optimal than *ND*^*i*^. Think of an *IGSD* strategy with only a non-zero cell differentiation probability from germ-like cells to soma-like cells and zero differentiation probabilities the other way around. Let’s assume it is the *i*th cell division that makes 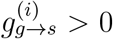, thus we have 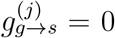 for *j* ≠ *i* and 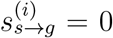 for any *i*. Based on Eq (12), the cell frequencies after the *n*th division are

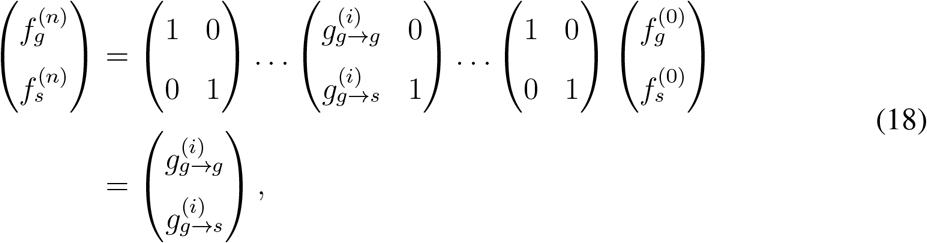

where 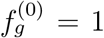and 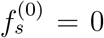. For convenience, we denote 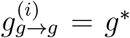 and 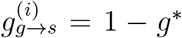.

From Eq (15), we have

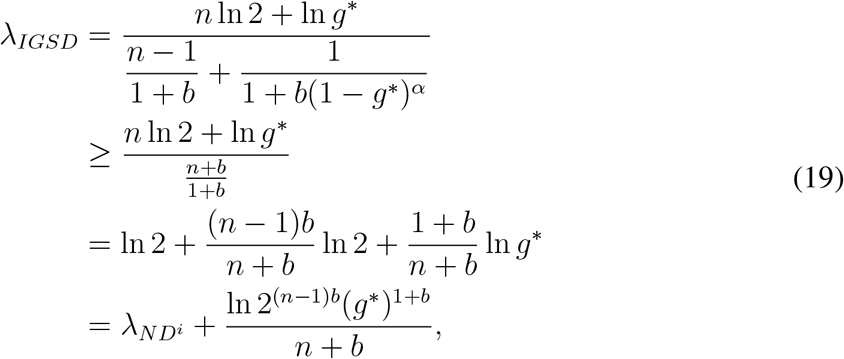

where we use (1 − *g*^∗^) ≥ 0 to obtain the inequality and 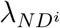 = ln 2 (appendix). Since *b* > 0 and *n* > 1, as long as 2^(*n*−1)*b*^ (*g*^∗^)^1+*b*^ ≥ 1, we obtain *λ*_*IGSD*_ > 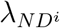. That is 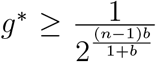 Therefore, when *b* > 0, we can always find an *IGSD* strategy with a 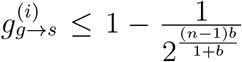 and all other 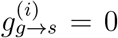 and *s* 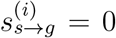, which leads to higher growth rate than *ND*^*i*^. Thus, *λ*_*IGSD*_ > 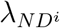. The proof of *λ*_*IGD*_ > *λ*_*IGSD*_ is in the appendix. Taken these together, we have *λ*_*IGD*_ > *λ*_*IGSD*_ > 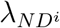.

### *RD* is optimal among *RD, ISD* and *ND*^*i*^

Next, we prove *λ*_*RD*_ > *λ*_*ISD*_ > 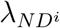. We first prove *λ*_*ISD*_ > 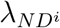. We prove that there exists an *ISD* strategy leading to a higher *λ* than 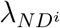 = ln 2 (appendix). Consider the *ISD* with 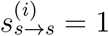, but with at least one *i* which makes 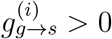i.e. 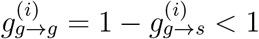 for 1 ≤ *i* ≤ *n*. The above constraint corresponds with the definition of the *ISD* strategy. Based on Eq (12), the cell frequencies after the *n*th division are

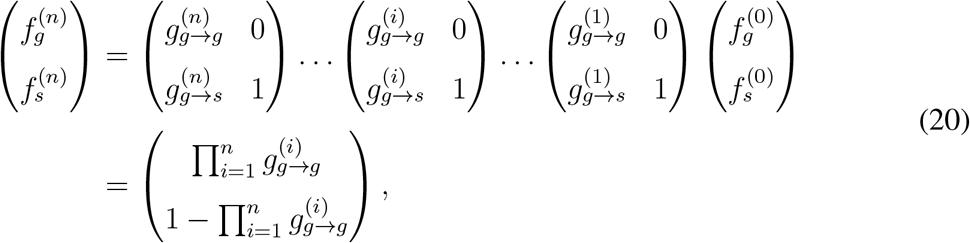

where *n* ≥ 1. Substituting Eq (20) into Eq (15) and together with *c* = 0, we find the growth rate of the ISD strategy.

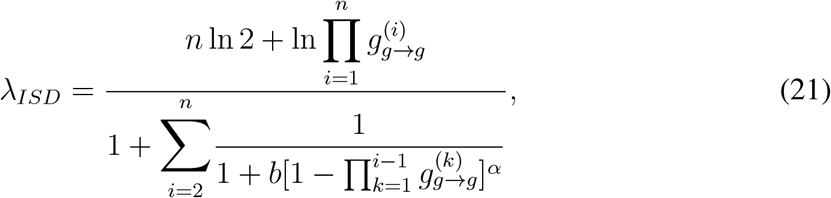

where the first item in the denominator represents the time for the first cell division *t*_1_ = 1 because of 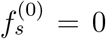. We define the second item of the denominator of Eq (21) as 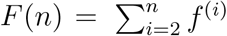, where 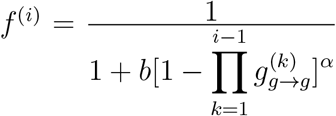. Next, we prove *F* (*n*) is a bounded function.

Since 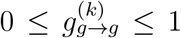, thus sequence 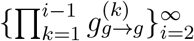 decreases with increasing *i*. That is, 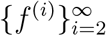 is a positive but decreasing sequence *f*^(2)^ is the largest one in 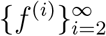. Therefore, the sequence 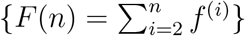 is an accelerating discrete sequence with respect to *n*. We have

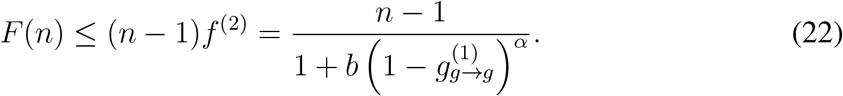

Substituting the right-hand side of inequality (22) into Eq (21), we have

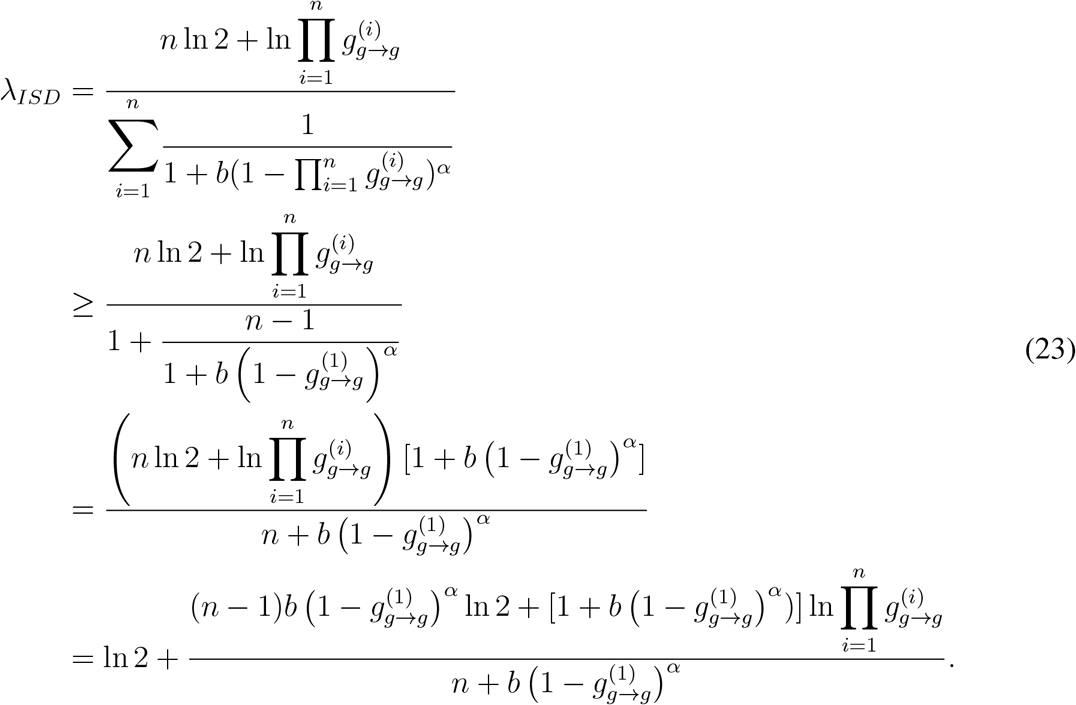

As long as there exist a strategy which makes the right side of Eq (23) greater than ln 2, we have 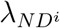 < *λ*_*ISD*_. Then we need to identify the conditions for

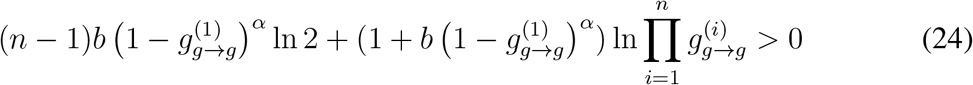

to hold. As *b* > 0 and *α* > 0, 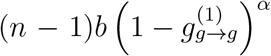 ln 2 ≥ 0. Since 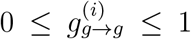, 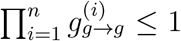 and ln 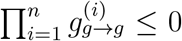. The second item of Eq (24) is negative. There exists a sequence 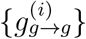, which makes 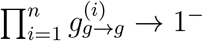 and

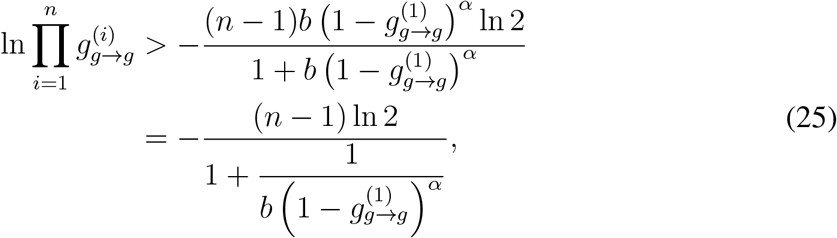

which makes the Eq (24) hold. With the above proof, we conclude that *λ*_*ISD*_ > 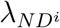 with only cell differentiation benefit. From Eq (25), we found that more *ISD* strategies are better than *ND*^*i*^ under high benefits *b*. However, when *b* is small, only *ISD* with 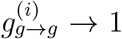 leads higher growth rate than *ND*^*i*^. The proof of *λ*_*RD*_ > *λ*_*ISD*_ can be found in the appendix. Thus, we have *λ*_*RD*_ > *λ*_*ISD*_ > 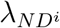. The results show that when there is a benefit and no costs, differentiation strategies (*ISD, IGSD, IGD* and *RD*) are better over *ND*^*i*^. Either *IGD* or *RD* is optimal under *b* > 0 and *c* = 0.

## S7 Appendix. *IGSD* and *ISD* cannot be optimal in the absence of either cell differentiation benefit or cost

In the appendix, we have proved *ND*^*i*^ is optimal in the absence of differentiation benefits, i.e. *b* = 0 and *c* > 0. Thus, we prove *IGSD* and *ISD* can be optimal in the absence of differentiation costs, i.e. *c* = 0 and *b* > 0. Since we also have proved that *ND*^*i*^ is optimal under *n* = 1 when *c* = 0 and *b* > 0 in appendix. Therefore, we only need to prove that the optimal strategy can neither be *IGSD* nor *ISD* when *b* > 0, *c* = 0 and *n* ≥ 2.

We first prove *λ*_*IGD*_ > *λ*_*IGSD*_. For a given *IGSD* strategy, we can always modify it and obtain an *IGD* strategy, which leads to a higher *λ* than the given *IGSD* strategy. For a given *IGSD* strategy, we know its transition probabilities 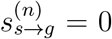. We modify the *IGSD* strategy by setting 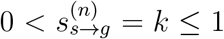 to get a *IGD* strategy. The constructed *IGD* strategy produces more offspring than the given *IGSD* strategy as its final number of germ-like cells is 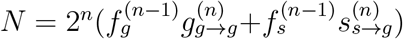, which is greater than that of the *IGSD* as 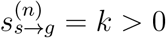 in the *IGD* strategy. Since there is no cell differentiation cost (*c* = 0), cell division rates are the same among all strategies. Thus, *λ*_*IGD*_ > *λ*_*IGSD*_.

Next, we prove *λ*_*RD*_ > *λ*_*ISD*_. Given an *ISD* strategy, we have 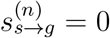. Construct a *RD* strategy that has the same transition probability matrixes as the given *ISD* strategy for the first (*n*−1) cell divisions. For the *n*th transition probability matrix, we keep the 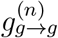 and 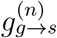 the same as that in the given *ISD* strategy. However, we set 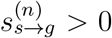 rather than 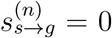 as that in *ISD* strategy. 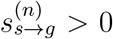 implies 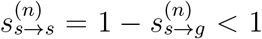. Then, *ISD* and *RD* have the same germ-like cells during the first *n* − 1 cell divisions. The fraction of germ-like cells for the *ISD* strategy after the *n*th cell divisions is 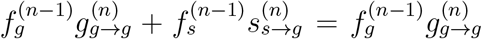 as 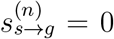 Whereas, the fraction of germ-like cells for the *RD* strategy after the *n*th cell divisions is 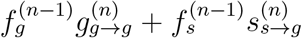.. Thus, the constructed *RD* has an extra 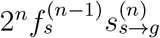. germ-like cells compared with the *ISD* strategy. Since the cell division rate at the *n*th cell division depends on the fraction of soma-like cells at the (*n* − 1)th cell division, the cell division rates *r*^(*i*)^ for the two strategies are the same, 1 ≤ *i* ≤ *n*. Thus, from Eq (15), we have

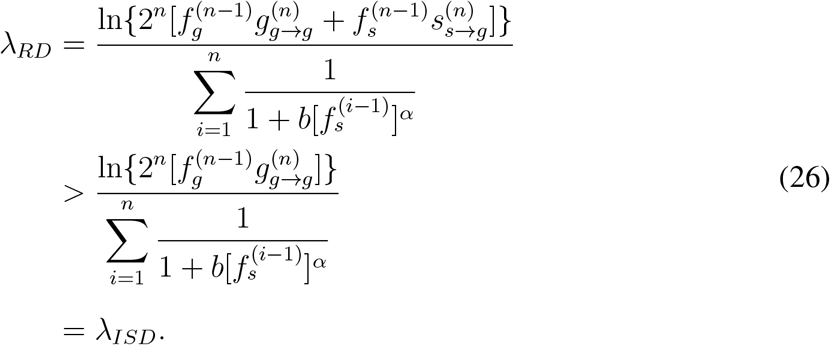

Therefore, *λ*_*RD*_ > *λ*_*ISD*_.

## S8 Appendix. Stage-dependent differentiation promotes irreversible cell differentiation under the effects of benefit function forms *α* and the ratio of differentiation costs between germ-like cells and soma-like cells *β*

Under the effects of *α* and *β*, we found that stage-dependent differentiation favors irreversible cell differentiation over stage-independent cell differentiation. *IGD* replaces stage-independent *RD* when *α* and *β* are both small, see Fig 8. Under this scenario, the cell transition probability *s*_*s*→*g*_ has a smaller effect in decreasing the growth rate than the transition probability *g*_*g*→*s*_. Thus, *IGD* produces a higher fraction of germ-like cells and bears less cell differentiation costs, leading to a higher growth rate. When *α* is around 1, *IGD* leads to faster growth than *ND*^*i*^. The reason is analogous to the one given in the main text.

**Figure 8:**
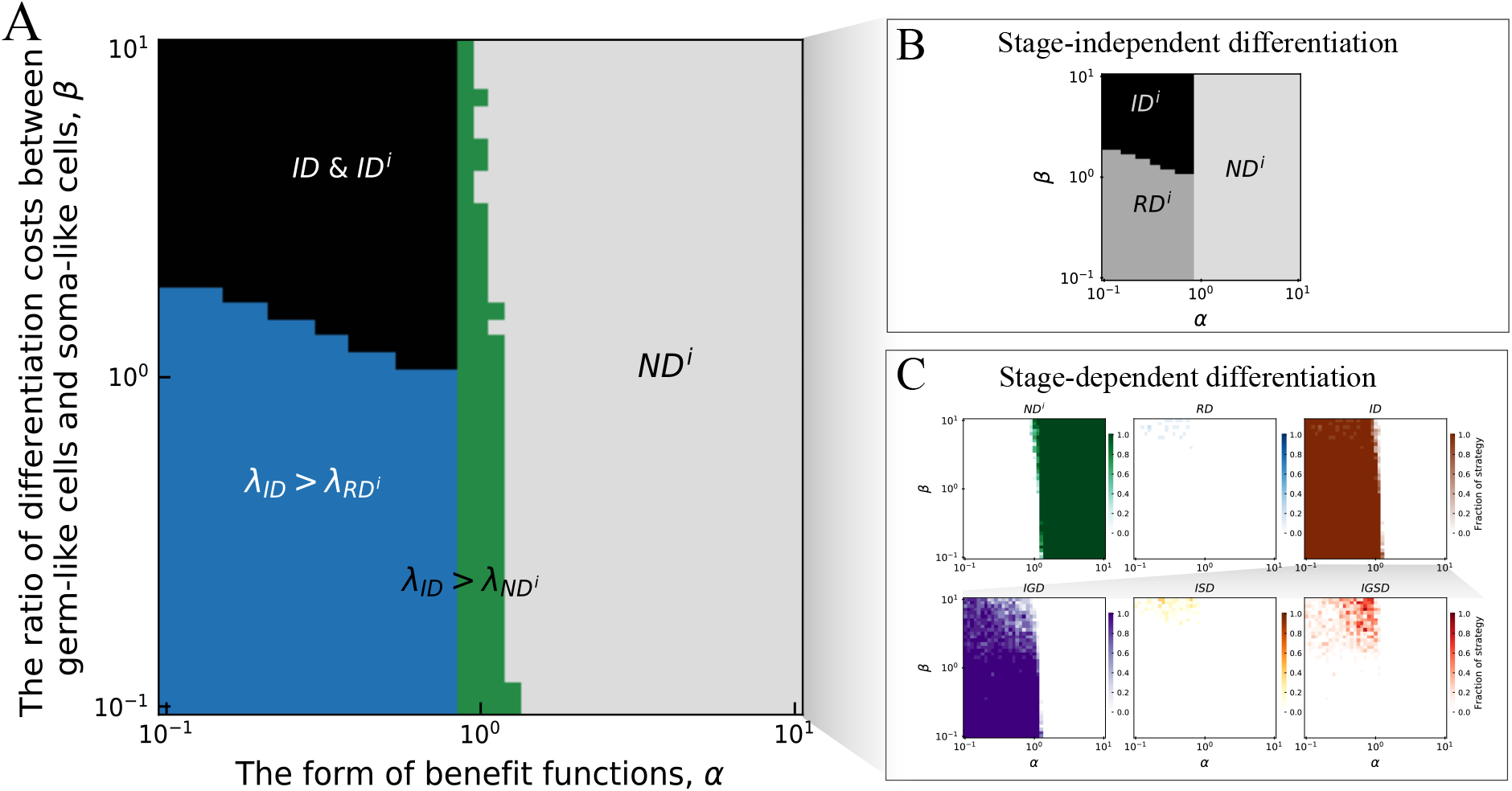
The effects of *α* and *β* on the growth rates of cell differentiation strategies. **A**. Comparison of the optimal strategy evolved in stage-independent and stage-dependent differentiation strategies depending on *α* and *β*. The areas of grey and black represent the parameter space in which the same strategy are optimal both under stage-independent and stage-dependent cell differentiation. The green area represents stage-dependent *ID* leading to a larger growth rate than stage-independent *ND*^*i*^. The blue strip represents stage-dependent *ID* leading to a larger growth rate than stage-independent *RD*^*i*^. **B**. The parameter space of optimal stage-independent differentiation strategy at different values of *α* and *β*. **C**. The frequencies of each stage-dependent strategy depending on *α* and *β*. Parameters for all panels *δ* = 0.1, *n* = 5 and *b* = *c* = 1. For calculating the growth rate of each strategy, see the appendix.

